# Mobius Assembly for Plant Systems highlights promoter-terminator interaction in gene regulation

**DOI:** 10.1101/2021.03.31.437819

**Authors:** Andreas I. Andreou, Jessica Nirkko, Marisol Ochoa-Villarreal, Naomi Nakayama

**Author notes:** Department of Bioengineering, Imperial College London (Naomi Nakayama).

## Abstract

Plant synthetic biology is a fast-evolving field that employs engineering principles to empower research and bioproduction in plant systems. Nevertheless, in the whole synthetic biology landscape, plant systems lag compared to microbial and mammalian systems. When it comes to multigene delivery to plants, the predictability of the outcome is decreased since it depends on three different chassis: *E. coli*, *Agrobacterium*, and the plant species. Here we aimed to develop standardised and streamlined tools for genetic engineering in plant synthetic biology. We have devised Mobius Assembly for Plant Systems (MAPS), a user-friendly Golden Gate Assembly system for fast and easy generation of complex DNA constructs. MAPS is based on a new group of small plant binary vectors (pMAPs) that contains an origin of replication from a cryptic plasmid of *Paracoccus pantotrophus*. The functionality of the pMAP vectors was confirmed by transforming the MM1 cell culture, demonstrating for the first time that plant transformation is dependent on the *Agrobacterium* strains and plasmids; plasmid stability was highly dependent on the plasmid and bacterial strain. We made a library of new short promoters and terminators and characterised them using a high-throughput protoplast expression assay. Our results underscored the strong influence of terminators in gene expression, and they altered the strength of promoters in some combinations and indicated the presence of synergistic interactions between promoters and terminators. Overall this work will further facilitate plant synthetic biology and contribute to improving its predictability, which is challenged by combinatorial interactions among the genetic parts, vectors, and chassis.

## INTRODUCTION

Synthetic biology focuses on design-led rational engineering of biological functions (1). When starting to establish a new chassis, tools and methods need to be established to enable predictive genetic engineering. The first step is to create and characterise reliably consistent standardised parts and vectors to control target gene expression. Multigene construction is necessary to engineer complex genetic circuits and regulatory networks underlying metabolic biosynthesis, developmental pathways, and environmental responses (2–5). An example is the transfer of nine genes in a single construct encoding fatty acid desaturases and elongases into the Indian mustard plant (*Brassica juncea*) to increase the yield of long-chain polyunsaturated fatty acids (6). Multigene transformation requires technologies for the successful generation of the constructs and their delivery. Design-led, precision molecular plant breeding for bioproduction of food, energy, materials, and pharmaceuticals has a great potential to catalyse a sustainable future (7, 8); however, the synthetic biology tools are still limited for plant systems. To enable complex multigene engineering of plant systems, here we report the development of suites of new standardised parts for gene regulation. The characterisation of these new parts has underlined often overlooked modes of gene regulation, which actually cast strong influences and so should be fully considered in genetic engineering and the research aiming to understand gene regulation mechanisms.

The generation of construct libraries of complex biosynthetic pathways demands a fast and easy DNA assembly method. We have developed a user-friendly and straightforward DNA assembly method, which combines high cloning capacity with a compact vector toolkit called Mobius Assembly (9, 10). Mobius Assembly employs IIS restriction endonucleases, which cleave DNA outside their recognition sequence allowing the unidirectional fusion of DNA sequences in a one-tube reaction (11, 12). The popularity of the type IIS methods has accelerated the development of new variants and improvements. It is now possible to predict the fidelity of the three or four base overhang ligations (13–15). At the same time, new methods have been developed to deal with domestication requirements (16), scars at coding sequence borders (17), the polarity of the transcriptional unit (TU) growth (18), backbone modularity and interoperability of DNA assembly (19, 20). Also, already established methods were adjusted for other organisms such as cyanobacteria (21, 22), fungi (23, 24) and Chlamydomonas (22). Additionally, the philosophy of universality in DNA assembly has gained further popularity(25, 26). Recent developments in the plant field include Loop Assembly (27) and an extension of the MoClo part toolkit for plants (28).

Plant binary vectors mediate the delivery and incorporation of transgene constructs into the genome of plant cells. Although functional, plant binary vectors could be improved further to enhance plant transformation efficiency. A typical plant vector for *Agrobacterium*-mediated transformation is comprised of left and right T-DNA borders for the transfer of the DNA construct into the host plant’s nuclear DNA genome (29); plasmid replication functions in *E. coli and Agrobacterium*; and markers for the selection and maintenance in *E. coli and Agrobacterium*. Currently, there are five categories of the vectors depending on the origin of replication in Agrobacterium: 1) RK2 (e.g. pCB series (30)) or RK2 in combination with ColE1 (e.g. pORE (31)); 2) pVS1 + ColE1 (e.g. pCambia (https://cambia.org)); 3) pSa with ColE1 (e.g. pGreen (32)); 4) pRi in with ColE1 (e.g. pCGN (33)), or F factor (eg pBIBAC (34)) or Phage P1 (eg. pYLTAC (35)); 5) BBR1 with pLX vectors (36). The pLX vectors revolutionised binary vector construction as they shifted from the cut-and-paste strategy to a modular design with minimal functional parts and introduction of stability features. Plasmid instability issues are among the main challenges of recombinant DNA production, affecting plasmid yield and quality (37). For plant binary vectors, the challenge is doubled since they should be stable in two different chassis, *E. coli* and Agrobacteria. The plasmid size (38), plasmid copy number (39), direct repeats (40), and inverted repeats (41), among others (37), are the factors affecting structural plasmid stability. Some popular plant vectors (e.g. pCambia) are based on bulky plasmid backbones (6, 2 kb), and thus they burden the size of the final constructs by default. Efforts have been made to generate binary backbones with a smaller size to prevent plasmid instabilities, such as pGreen (2.5 kb, (32)), pLSU (4.6 kb (42)), and pLX (3.3 kb (36)).

Stable transformation of whole plants is the common goal in plant genetic engineering, yet it is labour- and time-consuming. Transient assays by Agrobacterium-mediated DNA transfer in plant tissues (e.g. leaves) or transformation of isolated protoplasts directly with DNA are faster alternative systems (43). However, these need cultivated plants and are still preparation-intensive when it comes to testing libraries of constructs. Plant cell cultures combine the benefits of plant systems with those of microbial cultures, and they can be established as a next-generation synthetic biology chassis. They can be utilised for transient experiments or, with further improvement, for stable transgene integrations. Nonetheless, the methods and the part toolbox available for genetic engineering of plant cell cultures are as yet limited, and more characterisation and standardisation are wanted.

Promoters are DNA sequences near the transcription initiation site (TSS) responsible for the commencement and spatiotemporal gene transcription regulation, well known for their gene expression role. Terminators are genetic elements, usually located at the end of a gene or operon, and they terminate transcription. Although often underestimated, the terminator is likely to play important gene expression roles by controlling transcription arrest and mRNA stability and tuning other transcription functions. Currently, the most commonly used promoters and terminators in plant systems come from the cauliflower mosaic virus (*35S*), Agrobacterium opine genes (*NOS, MAS, OCS*), and recently from the plant genome (*UBQ10pro* and *HSPter* from Arabidopsis). The shortage of well-characterised genetic elements results in the repeated use of the same parts within a multigene construct. This might negatively influence plasmid stability due to repetitive sequences (40) or even raise the frequencies of homologous-dependent gene silencing (44). Consequently, a plant genetic engineering toolbox is wanting more promoters and terminators. Besides, there is a need for short parts, especially crucial for single vector multigene delivery; they will substantially reduce the already large size of multi-TU constructs. Large constructs tend to make plasmids structurally unstable (38) and decrease the efficiency of agrobacterium-mediated transformation in plants (45).

Here we report the development of Mobius Assembly for the engineering of plant chassis. Mobius Assembly for Plant Systems (MAPS) includes new plant binary vectors (pMAP) compatible with a wide variety of plant systems, such as plant cells, tissues, and whole organisms. pMAP vectors exploit the WKS1 origin of replication from a *Paracoccus pantotrophus* cryptic plasmid. They are small in size, structurally stable and suitable for transformation methods that demand high yield plasmid DNA. As proof of concept, we tested the functionality of pMAP vectors on the MM1 plant cell line with great success. For the first time, we demonstrated that plant cell culture transformation is Agrobacterium strain and plasmid specific, paving the way to establish plant cell cultures as a next-generation synthetic biology chassis. We also showed that plasmid stability is strain and vector-specific, and we stressed the importance of testing plasmid stability before cloning experiments. MAPS is also equipped with a selection of new standardised parts (Phytobricks) of short promoters and terminators, reducing the overall construct size and driving a gradient of gene expression levels.

Throughout the characterisation experiments, we demonstrated the importance of the terminators in regulating gene expression. We also revealed a synergistic effect between the terminators and the promoters, which has a vital role in tuning the expression.

## MATERIALS AND METHODS

### Bacterial strains and growth conditions

*E. coli* strains DH5α, DH10B, TOP10, JM109, NEB stable, and *A. tumefaciens* strains AGL-1, C58C1*, GV3101, LBA4404 were used. *E. coli* chemically competent cells were made using the TSS preparation, described by Chung & Miller (46), except TOP10, which were bought from ThermoFisher Scientific. *A. tumefaciens* electrocompetent cells were prepared as follows: Overnight seed culture was grown in S.O.C medium (0.5 ml) at 28^°^C, 200 min^-1^ with the appropriate antibiotics, was diluted in 500 ml of the same fresh medium. After overnight incubation at the same conditions, and when the culture reached OD_600_ 0.6-1.0, cells were harvested and washed three times with ice-cold 10% sterile glycerol. Finally, cells were resuspended in 1.5 ml 10% cold glycerol, aliquoted in 40 μl batches and snap-frozen in liquid nitrogen. *E. coli* cells were incubated at 37°C (30°C for NEB stable), 200 *min^-1^* either in 5 ml (for a high copy) or 10 ml (for a low copy), or 100 ml (for midi prep) in LB growth medium supplemented with antibiotics. Cells bearing the *LhGR-pOp6* and *sXVE-lexA* inducible systems were grown for 24 h instead due to their slower growth rate. Agrobacterium cells were cultured in 10 ml YEP medium at 28°C, 200 *min^-1,^* for two days.

### Plasmid isolation

Plasmids were isolated using Monarch (NEB) or PureYield (Promega) or GeneJET (Thermo Fisher Scientific) Plasmid Miniprep Kits. The Promega PureYield Plasmid Midiprep System (Promega) with vacuum manifold and GeneJET Plasmid Midiprep Kit (Thermo Fisher Scientific) were used for higher yields. Plasmid isolation from Agrobacteria was carried out with 1.5 times the recommended reagent volumes.

### Bacterial transformations

For *E. coli* transformation, 5 μl of the plasmid DNA was incubated with 100 μl of the competent cells (25 μl for TOP10) on ice for 30 min, followed by a heat shock at 42°C for 90 sec (30 sec for TOP10) and re-cooled on ice for 5 min. S.O.C medium (400 μl) was added, and after 1 h incubation at 37°C, 100 μl of the cell suspension was plated on LB agar plates with antibiotic selection. For Agrobacterium transformation, 1 μl of plasmid DNA was added into an aliquot of competent cells and incubated on ice for 5 min. Subsequently, the cells were transferred into an ice-cold electroporation cuvette, 0.2 cm gap (Bio-Rad) and pulsed twice in a Gene Pulser Xcell Electroporation System (Bio-Rad) using pre-set settings for Agrobacteria (25 μF, 200 Ω, 2400 V, 2 mm). S.O.C medium was added immediately (1 ml), and after 2h of shaking at room temperature, 20 μl was plated on YEP plates with the correct antibiotics and incubated at 30°C for 2-3 days until colonies had formed.

### Molecular cloning

All PCR amplifications for plasmid construction and cloning were performed using Q5® High-Fidelity DNA Polymerase (NEB), followed by purification with Monarch® PCR & DNA Cleanup Kit (NEB). The Mobius Assemblies were verified first by colony PCR using GoTaq® Green Master Mix (Promega) and then with double restriction digestion with *EcoRI*-HF (NEB) and *PstI*-HF (NEB). The constructs were further verified by Sanger sequencing (GATC Biotech-Eurofins or Edinburgh Genomics).

### Plasmid construction

Mobius Assembly Level 1 and Level 2, Acceptor Vectors for plants, were built using Gibson Assembly. The Mobius Assembly cassettes were amplified from the corresponding *E. coli* plasmids in the Mobius Assembly Vector toolkit and fused to the plant binary vectors. pGreen based Level 1 vectors were constructed using pGreen0029 (32), which was acquired from the John Innes Centre. NptI gene was replaced with a spectinomycin resistance gene amplified from pCR8 vector (ThermoFisher Scientific) to generate Level 2 vectors. pLX based vectors Level 1 and Level 2 were built using pLX-B3 and pLX-B2 (36), respectively, which were acquired from Centro Nacional de Biotecnología (CNB-CSIC).

WKS1 Ori was synthesised by Twist Bioscience into two parts (due to repetitive sequences), and pUC Ori was amplified from pUC19, both flanked by *BsaI* recognition sites. The minimum sequence requirement of pUC for stable replication in *E. coli* was found to include RNA I/RNA II transcripts on 5′-side and dnaA/dnaA′ boxes on the 3′-side, while co-directional transcription of two different replicons in the same plasmid was shown to increase transformation efficiency and DNA yield (42) A pLX Level 1 A vector with the construct *NOS:BglR:NOS-UBQ10:nluc:HSP* was amplified outside the BBR1 Ori with primers harbouring *BsaI* recognition sites and fused with pUC and WKS1 Ori. The resulting plasmid was used as a template to amplify the pUC-WKS1 fused Ori, which then replaced BBR1 Ori from pLX Level 1A and Level 2A vectors with Gibson Assembly, resulting in the pMAP Level 1A and Level 2A vectors. The rest of the pMAP vectors were constructed again using isothermal assembly and as a template for the Mobius cloning cassettes, the plasmids from the original Mobius Assembly kit (9). MethylAble modules were devised with the Gibson Assembly using mUAV as a template. Overlapping primers bearing two outward-facing *BsaI* sites prone to CpG methylation and the suitable standard overhangs were used to amplify mUAV into two parts. The resulting parts were purified, digested with *DpnI* (*ThermoFisher Scientific) to* eliminate the template DNA, and they were fused in an isothermal reaction.

### Mobius Assembly cloning

A detailed protocol can be found in (10). Briefly, the assembly was performed in a one-tube reaction with a total volume of 10 μl, with ∼50 ng Acceptor Vectors and double amounts of inserts. Reagents added were 1 μl of 1 mg/ml BSA (diluted from 20 mg/ml - NEB), 1 μl T4 DNA ligase buffer (NEB or Thermo Fisher Scientific), 0.5 μl *AarI* (ThermoFisher Scientific) and 0.2 μl 50x oligos (*AarI* recognition site) for Level 0 and Level 2 cloning or *Eco31I/BsaI-HFv2* (ThermoFisher Scientific/NEB) for Level 1 cloning, and 0.5 μl T4 DNA ligase (NEB or Thermo Fisher Scientific). The reactions were incubated in a thermocycler for 5-10 cycles of 5 min at 37°C and 10 min at 16°C, followed by 5 min digestion at 37°C and 5 min deactivation at 80°C. (5 cycles for Level 0 and the first round of Level 1 cloning – 10 cycles for Level 2 and large constructs >10 kb).

### CpG Methylation

Midi-prep plasmid DNA of the MethylAble modules was used. The reaction was carried out in 20 μl total volume according to the NEB protocol. Briefly, up to 2 μg of plasmid DNA was incubated for 4 h at 37°C with 2 μl of CpG Methyltransferase from *M.SssI* (NEB) in 1× Methyltransferase Reaction Buffer supplemented with 2 μl of diluted SAM (6.4 mM). The reaction was stopped by heating to 65°C for 20 min and was purified by column chromatography.

### Selection and amplification of Arabidopsis promoters and terminators

Ubiquitously and likely constitutively expressed genes used in qPCR or genes expressed in the plant embryos were identified through literature research. The most stably expressed were selected for promoter and terminator identification. Their sequences were retrieved from the TAIR webpage (47) (http://www.arabidopsis.org/index.jsp) and blasted in NCBI to find the untranslated regions flanking the genes. A 1.5 kb sequence upstream of the start codon was run through the online prediction software and TSSPlant (48) (http://www.softberry.com) to identify TATA and TATA-less promoters or enhancer sites. In the promoter selection, it was also considered, when possible, to include the initiator (INR) elements (YYA(+1)NWYY-TYA(+1)YYN-TYA(+1)GGG)) and downstream promoter (DPE) element (RGWYV). Finally, two promoter versions were created based on the sequence size, one ∼300 bp and one ∼500 bp. Potential promoter elements linked to increased gene expression were identified using PlantCare (49) and PLACE (50) software.

Terminators were selected with PASPA, a web server for poly(A) site prediction in plants and algae (51) (http://bmi.xmu.edu.cn/paspa/interface/run_PASPA.php). A sequence 300bp downstream of the stop codon was run into PASPA, and the selection of the terminator was set 10bp after the second polyadenylation site, resulting in a sequence of around 200bp. They were then analysed in RegRNA2.0 to identify other RNA functional motifs (52) (http://regrna2.mbc.nctu.edu.tw). Appropriate primers were designed for both promoters and terminators for cloning into mUAV.

### Functional dissection of FAD2 and HSP terminators

Primers were designed to gradually remove functional sequence elements from the 3’ end of each terminator through PCR and subsequently cloned to mUAV. Site-directed mutagenesis by Gibson Assembly was employed to mutate the poly-A signal of the *HSP* part3, while the same method was used to build *HSP* part 5.

### Plant-cell suspension culture and Agrobacterium mediated transformation

MM1 cell suspension culture of *Arabidopsis thaliana* ecotype Landsberg *erecta* (53) was subcultured weekly by 1:10 dilution in 1× MS medium (Sigma) supplemented with 2% w/v glucose (Sigma), 0.5 mg/L NAA (Sigma), 0.05 mg/L kinetin (Sigma), with the pH adjusted to 5.8 with KOH. Agrobacterium-mediated cell culture transformations were performed according to a modified protocol (54) under sterile conditions. Freshly prepared Agrobacterium plates (2-3 day-old maximum) and Arabidopsis cells 5-7 days post-sub-culturing were used. 25 ml of cell culture was transferred to a 50 ml falcon tube and centrifuged at 100 min^-1^, at 22°C for 10 min. The supernatant was removed with a 25 ml serological pipette, and the cells were resuspended in 40 ml of transformation medium at pH 5.8 (1× MS plant salt mixture without NH_4_NO_3_ (Sigma-Aldrich), 1% w/v sucrose, 1% v/v B5 vitamin stock solution and 1 μg/ml 2, 4-dichlorophenoxyacetic acid (Sigma-Aldrich)) and 25 μl of 100 mg/mL freshly prepared acetosyringone (Sigma-Aldrich) was added. B5 vitamin stock solution was prepared with 0.4 g/l nicotinic acid (Sigma-Aldrich), 0.4 g/l pyridoxine-HCl (Supelco/Sigma-Aldrich), 4 g/l thiamine-HCl (Merck), and 40 g/l myo-inositol (Merck). After gentle resuspension, 4 ml of plant cells were transferred in each well of a 6-well plate, and inoculated with a small amount of agrobacterium directly from the petri dish with the help of a 200 μl pipette. After 2 days of incubation on a rotary shaker, plant cells were supplemented with 5 ml of fresh culture medium and 50 μl of cefotaxime (Arcos) at 100 mg/mL which was added to eliminate agrobacteria.

### Arabidopsis protoplast isolation and transformation

*Arabidopsis thaliana* (Wildtype Col-0) seeds were sown on the soil. After a 2-day pre-treatment at 4°C in darkness, they were grown under long-day conditions (21°C; 16h light / 8h dark cycles; light intensity ∼100μmol/m^2^s^-1^; humidity ∼65%) for two weeks. Then, seedlings were transplanted and moved to short-day conditions (21°C; 9h light / 15h dark cycles; light intensity ∼110μmol/m2s-1) and grown for 4-6 weeks until harvest.

The protoplast isolation and transformation protocol was developed based on (55) and (56). Briefly, leaves from 6-8 week old plants were digested with MGG digestion solution containing: Cellulase ONOZUKA R-10 (0.2%), MACEROZYME R-10 (0.06%) (Yakult Pharmaceutical) and Driselase (0.08%) (Sigma-Aldrich). The following day, the protoplasts were filtered and washed 3-5 times with MGG without enzymes and resuspended in MMM solution (0.4 M mannitol, 15 mM MgCl_2_, 0.1% w/v MES, pH 8) in a concentration of 5×10^5^ cells/ml. Protoplasts were transformed into 1 ml 96-well plates. 8 μl DNA (500 ng/μl) was added to 75 μl of the protoplast suspension, followed by addition of 83 μl PEG (0.4 M mannitol, 0.1 M Ca(NO_3_)_2_-4H_2_O, 40% PEG4000, pH8) and 1 min incubation. Subsequently, the tubes were filled with MGG solution and incubated for 1 h at RT. After the incubation, the solution was removed, and the protoplasts were resuspended in 100 μl of fresh MGG and incubated overnight in darkness at RT. For the inducible system, 0.1% EtOH, 2.5 μM DEX (Acros) or 5 μM β-estradiol (LKT laboratories) were added.

### Luciferase assay

The plates containing the transformed protoplasts were centrifuged at 200 min^-1^ (acc/decc = 3) for 10 min, and 60 μl of supernatant was discarded. The protoplasts were resuspended, and 40 μl was transferred to white optical plates in a grid pattern with empty spaces between wells to reduce luminescence bleed-through. For the MM1 transformed cells, 40 μl were transferred with a cut-tip directly from the 6/12-well plates to 96-well plates in triplicate. Luciferase activity was assayed in an Omega luminescence plate-reader (Fluostar) with four different gains following the instructions of the Nano Dual-Luciferase® Reporter kit (Promega). A further correction for luminescence bleed-through was applied using the software developed by Mauri *et al*. (57).

## RESULTS

### Mobius Assembly for Plant Systems (MAPS)

MAPS is an extension of our molecular cloning system Mobius Assembly (9) to mediate transformation and gene expression into plant systems, either whole-plant or cell-based. MAPS iterates between two cloning levels enabling quadruple augmentation of transcriptional units (**Figure 1**). The introduction of the rare cutter *AarI* minimises the need for removing internal restriction sites, and the exploitation of constitutively expressed chromogenic proteins for clonal screening eliminates the need for additives in the selection plates. MAPS vector toolkit consists of a core set of pMAP cloning/destination vectors (Level 1 Acceptor Vectors Α-Δ and Level 2 Acceptor Vectors Α-Δ), which have a fusion origin of replication (WKS1+pUC) to replicate in *E. coli* and Agrobacteria.

**Figure 1.**
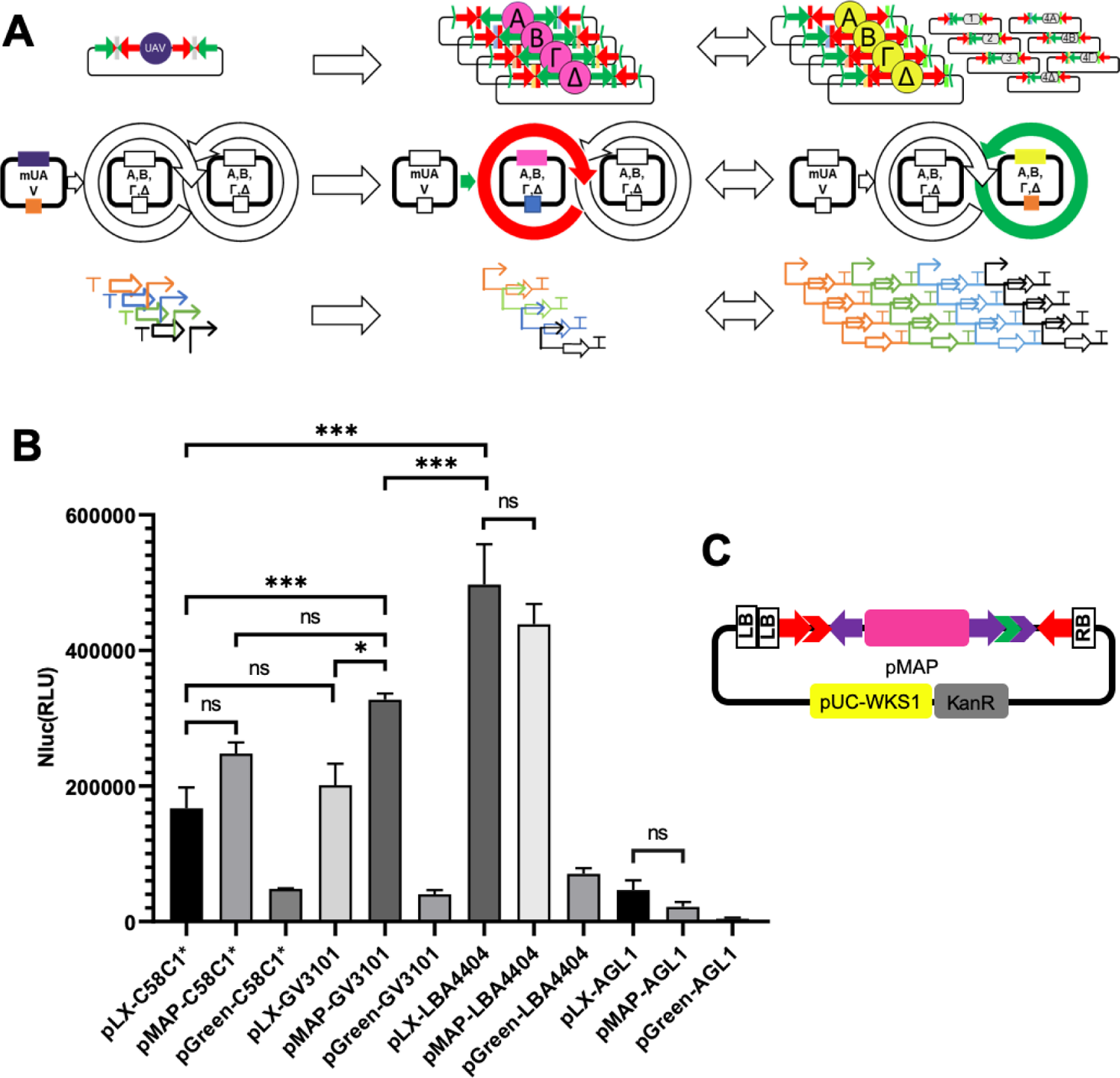
Mobius Assembly for Plant Systems (MAPS). **A.** Schematics of how Mobius Assembly operates. MAPS is an adaptation of the Mobius Assembly for plant systems based on small plant binary vectors. The core vector toolkit is comprised of one storage vector in Level 0 (mUAV), four pMAP Level 1 Acceptor Vectors, four pMAP Level 2 Acceptor Vectors and seven Auxiliary plasmids. It employs *BsaI* in Level 1 cloning and the rare cutter *AarI* in Level 0 and Level 2 cloning. **B.** Proof of concept transformation of MAPS vectors in the MM1 Arabidopsis cell line. The construct *NOS:BglR:NOS-UBQ10:nluc:HSP* was cloned either in pLX, pGreen and pMAP Level 1 Vector, and it was transformed in C58C1*, GV3101, LBA4404 and AGL-1 Agrobacterium strains. The Agrobacteria bearing the binary vectors were co-cultured with cells from the MM1 cell line in 6-well plates. After two days, Agrobacteria were eliminated with Cefotaxime. Two more days later, the cells were assayed for Nano luciferase activity in a plate reader. **C.** Plasmid design of pMAP. Nluc= Nano luciferase activity, bar graphs show luciferase activity values in mean±SE, p < 0.05, one-way ANOVA and Tukey’s HSD test.

Initially, we used pGreen based vectors for the transient transformation of Arabidopsis mesophyll protoplasts; nevertheless, we found to have transformation issues with large constructs and proved unsuitable for transforming plant cell cultures. Also, it was reported to have instability issues (58). Therefore, we built a new small plant binary vector based on the pLX architecture, called pMAP (**Figure 1C**), suitable for protoplast, cell culture, tobacco leaves, and whole-plant stable transformation. To achieve this, we devised a new origin of replication for plant binary vectors comprised of the fusion of pWKS1 and pUC19 Ori. pWKS1 Ori derives from a small, multicopy and cryptic plasmid pWKS1 (2697 bp) of *Paracoccus pantotrophus* DSM 11072, which has only replication and mobility function domains (59). Cryptic plasmids usually encode proteins involved in plasmid replication and mobilisation, and they do not have obvious benefits to the host cells that carry them (60, 61). The replication domain consists of an origin of vegetative replication OriV (249 bp) and the gene encoding its cognate binding protein RepA (1020 bp). pWKS1 Ori was found to be functional in *Agrobacterium tumefaciens* but not in *E. coli* (59). The minimal stable pUC Ori from pUC19 was amplified and fused in co-directional transcriptional orientation with pWKS1 Ori. The new origin of replication is only 37.84% larger than BBR1 (1978 bp instead of 1435 bp), which is still smaller than the short version of pVS1 + colE1 in pLSU (2654 bp, (62) and pSa+colE1 in pGreen (2179 bp without the plasmid stability domains, (32).

The kit also has pLX (BBR1 and RK2) based destination vectors (Level 1 and Level 2 Acceptor Vectors A), which are medium and low copy number vectors, respectively, to deal with possible instabilities that occur in large constructs housing repetitive or similar sequences. The mUAV and the seven Auxiliary plasmids are also included, as described in the original Mobius Assembly kit (9). The MAPS toolkit also contains a selection of plant promoters, terminators, antibiotic resistance genes, and reporter genes. All MAPS plasmids are listed in **Supplementary Table S1** and available through AddGene (https://www.addgene.org/browse/article/28211394/).

To test the MAPS vector functions, we transformed the different variations of MAPS vectors (pGreen, pLX or pMAP) to transform Arabidopsis cell suspension culture MM1 (63) into four widely used Agrobacterium strains (GV3101, AGL1, C58C1*, LBA4404 (**Supplementary Table 2**, **Figure 1B, C**). The evaluation was carried out by a luciferase assay using the construct *NOS:BlpR:NOS-UBQ10:nluc:HSP*. The results showed that all the Agrobacteria strains could infect the MM1 cell line (**Figure 1B**). The LBA4404 strain exhibited the highest transformation efficiency giving the most top luciferase signal. The second-best activity was found in the GV3101 and C58C1* strains. GV3101 was more efficient in transfection than C58C1*, harbouring the pMAP vector, while there was no significant difference when bearing the pLX vector. On the other hand, the AGL1 strain had the lowest transfection capacity for all the binary vectors tested. Regarding the binary vectors, pMAP performed better than pLX in the GV3101 strain, leading to high luciferase values, while there was no statistical difference with C58C1*, LBA4404 and AGL1 strains. Strikingly, pGreen performed poorly in all Agrobacteria strains compared to the other vectors. All the vectors have been tested for their stability in both *E. coli*, and Agrobacterium strains exhibiting different stabilities in different strains (**Supplementary results**).

### MethylAble feature

Combinatorial DNA library is a powerful method for applications such as part characterisation and metabolic pathway optimisation (64). However, manual generation of a combinatorial DNA library takes time, effort and resources, especially when it involves several cloning steps. Therefore, we developed a new feature called ‘MethylAble’ to propagate intact *BsaI* recognition sites by cytosine methylation during Level 1 cloning so that Level 2 constructs can directly receive Phytobricks (**Figure 2**). According to Rebase (http://rebase.neb.com), GGTCTC^m5^/C_m5_CAGAG methylation protects *BsaI* digestion. An *amilCP* expression cassette was designed to carry in each site divergent and convergent *BsaI* recognition sites bordering the four base pair overhangs. The overhangs correspond to the part that the MethylAble replaces. The divergent *BsaI* recognition sites were designed to be prone to CpG methylation (CGGTCTC^m5^G/GC^m5^CAGAGC), and consequently, *BsaI* digestion is blocked, while the convergent sites (TGGTCTC^m5^T/AC^m5^CAGAGA) are not.

**Figure 2.**
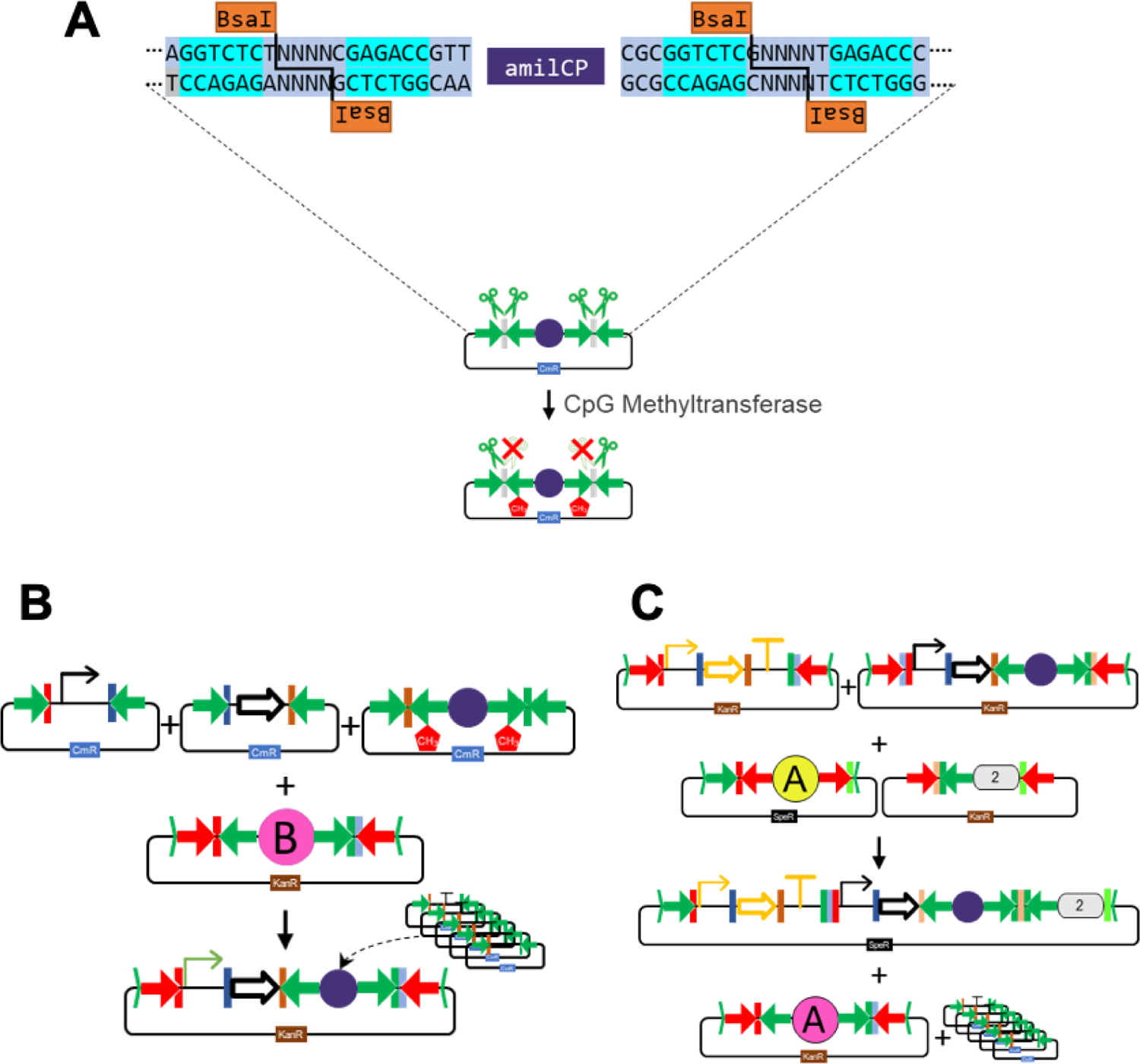
MethylAble feature for transcending levels in Mobius Assembly. MethyAble is a new feature of the Mobius Assembly that enables direct assembly of standard parts with Level 2 constructs. **A.** MethylAble plasmid: It has an *amilCP* gene flanked by inward and outward-facing *BsaI* recognition sites, which both cleave at the Phytobrick overhangs. The outward-facing *BsaI* sites are blocked with CpG methylation. **B.** In this example, the MethylAble plasmid replaces the terminator part, which is fused with a promoter and a coding sequence in a Level 1 reaction. **C.** If desired, the Level 1 construct can directly receive Phytobricks. The Level 1 construct is then fused with a second TU, resulting in a 2-TU construct. This construct can be fused directly with a library of Phytobricks in a Level 1 reaction. Pink circles demarcate Level 1 and yellow circles Level 2 Acceptor Vectors, respectively; the purple circle shows the MethylAble cassette and oval numbered shape the Auxiliary Plasmid. The green and red arrows are *BsaI* and *AarI*, respectively.

Upon CpG methylation, the *amilCP* TU is cloned with intact divergent *BsaI* sites in a Level 1 reaction and subsequently to a Level 2 reaction (**Figure 2**). Therefore, Level 0 parts can directly be inserted in a premade Level 1 TU or fused to a Level 2 TU. As *amilCP* chromoprotein will be expressed, the correct constructs will have a purple colour until the Level 0 parts replace the *amilCP* expression cassette. The MethylAble feature can be designed for any standard part, or combinations of parts, by setting the appropriate overhangs. It is also possible to use this feature to change the polarity of the TU growth in a Level 2 reaction by replacing the standard overhangs with the Level 1 overhangs. MethylAble plasmids are built with isothermal assembly using mUAV as a template and overlapping primers to introduce the *BsaI* recognition sites and the selected overhangs.

As a proof of concept, the MethylAble Feature was used to build the library of the three inducible promoters, each of which was combined with 14 new terminator parts. The MethylaAble cassette was designed to have the terminator overhangs (GCTT-CGCT). In Level 1, the constructs were made, as shown in (**Supplementary Table S3**). Briefly, a normaliser unit was built in Vector A, the three trans-activator units in Vector B and three inducible promoters were combined with the MethylAble feature in Vector Γ. Without the MethylAble feature, the inducible promoters should have been combined with all terminators in this step. Then in a Level 2 reaction, the normaliser, the trans-activators and the inducible promoters were fused to generate three constructs in total. Lastly, in a Level 1 reaction, the inducible constructs were combined with Level 0 terminators to give the final 14 constructs.

### Design and characterisation of new promoters and terminators

To select new promoter and terminator parts, we mainly opted for likely ubiquitously expressed genes (**Supplementary Table S4**), which have a good chance of driving high expression consistently in different applications for plant research. Ubiquitously expressed genes used as positive controls in qPCR were identified through literature search (65), and the most stably expressed were selected for promoter and terminator identification. *LEC2* promoter was selected for expression in undifferentiated cells, similar to cells in suspension cultures (66). The first screening round included short promoters (∼300bp) and short terminators (∼200bp) from the genes *ACT2*, *FAD2*, *TUB9*, *APT1*, *NDUFA8* and *LEC2*. As the ∼300 bp promoters’ activity was low (Data not shown), we designed new promoters derived from the genes *TUB2*, *UBQ11*, *UBQ4*, *ACT7*, and the longer versions of the previous promoters (∼500 kb).

The library of the short promoters and terminators was characterised by the high-throughput protocol for transient gene expression assay with Arabidopsis mesophyll protoplast in 96-well format, which we developed based on (56, 67). Specific parameters were optimised to improve the transformation efficiency and reproducibility, simplify the protocol and reduce the reagents. Nano-Glo Dual-Luciferase reporter assay system was used for the characterisation experiments, as it has better sensitivity at low expression levels and improved reagent stability. Nano luciferase was compatible with established instrumentation and protocols for firefly luciferase and suitable for plant expression (68). The Nano luciferase gene was employed as a primary reporter, and its signal was normalised to a construct bearing the firefly luciferase flanked by *UBQ10* promoter and *UBQ5* terminator to minimise the expression variations.

For the evaluation of the promoters, the *NLuc* transcription was terminated either by *NDUFA8* or *HSP* terminator, while 17 different promoters drove the transcription initiation. *UBQ10* exhibited by far the highest expression activity among the promoters, followed by *MAS* when combined either with *HSP* or *NDUFA8* terminator (**Figure 3A and B**). With the exemption of the *TUB9* promoter, the *HSP* terminator resulted in more elevated expression than *NDUFA8* to the various promoters. Two of the new isolated promoters, *UBQ11* and *UBQ4*, were found to have a comparable or even better expression from the commonly used *35S* and *OCS* promoters. *UBQ11* had an expression of 68.3 RLU with *HSP* and 21.8 RLU with *NDUFA8*, and *UBQ4* showed an expression of 66 RLU fused to *HSP* and 16 RLU combined with *NDUFA8*. The corresponding values of the 35S promoter were 41.7 RLU and 20.7 RLU, in combination with *HSP* and *NDUFA8* terminators, respectively. Furthermore, the newly isolated promoters *ACT7*, *TUB2*, *TUB9*, *APT1*, *ACT2* and *LEC2* drove stronger expression from the *NOS* promoter. Lastly, the lowest expression was found for *FAD2* and *NDUFA8* promoters.

**Figure 3.**
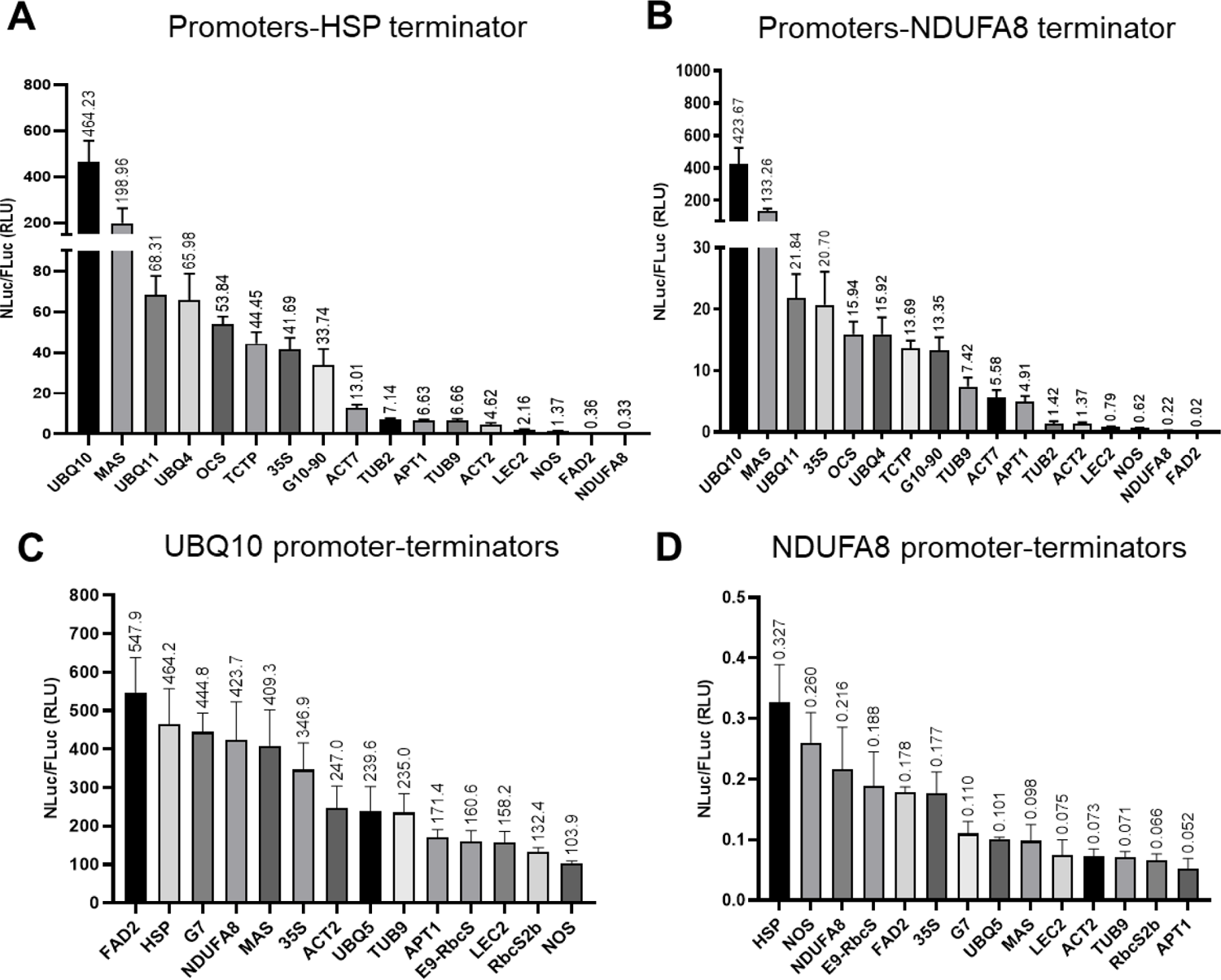
High-throughput characterisation of plant promoter and terminator part libraries. The characterisation was carried out with PEG-transformation of Arabidopsis leaf mesophyll protoplasts in 96-well plates. The expression strength of the promoters and terminators was assayed in 96-well plates in a plate reader measuring the Nano luciferase activity normalised by Firefly luciferase. The promoter library was characterised either with *HSP* (**A**) or *NDUFA8* (**B**) terminator, while the terminator library was characterised either with *UBQ10* (**C**) or *NDUFA8* (**D**) promoter. The construct for the promoter study was *Promoter:nluc:HSP/NDUFA8:UBQ10:fluc:UBQ5* and for the terminator characterisation *UBQ10/NDUFA8:nluc-Terminator:UBQ10:fluc:UBQ5*, both housed in pGreen based vector. Nluc= Nano and Fluc=Firefly luciferase activity, bar graphs show luciferase activity values in mean±SE.

For the terminators’ evaluation, the strong *UBQ10* or the weak *NDUFA8* promoter drove the NLuc expression, terminated by one of the 14 terminators. The different terminators resulted in a wide range of expression spanning 5.3 and 6.3 orders of magnitude for *UBQ10* and *NDUFA8* promoters, respectively (**Figure 3C and D**). For the strong *UBQ10* promoter, the *FAD2* terminator resulted in the highest activity with 547.9 RLU and NOS in the lowest with 103.9 RLU. For the weak *NDUFA8* promoter, the *HSP* terminator led to the highest expression with 0.327 RLU and *APT1* to the lowest at 0.052 RLU.

Since it was surprising to see a wide range of expression levels different terminators drive with the same promoters, we characterised them more with three chemical inducible systems commonly used in plant sciences: dexamethasone, estradiol and ethanol inducible systems (69). They are all based on two-component mechanisms involving at least two transcriptional units, in which the exogenously applied chemical activates the transcription factor that further ‘transactivate’ the downstream target genes. The target genes are driven by the specific promoters that contain binding sites for the transactivator (*pOp6*, *lexA* and *alcSynth*, respectively), and hence promoters cannot be changed. Terminators, on the other hand, can be altered. Interestingly, terminators combined with *pOp6-35S* and *lexA-35S* inducible promoters resulted in narrow and more uniform luciferase expression, spanning at around 2.2 and 2.9 orders of magnitude, respectively (**Figure 4A, B**). For *lexA-35S*, the expression is spread between 37.9 and 111.6 RLU when combined with *UBQ5* and *35S* terminators, respectively. For *pOp6-35S*, the expression ranges between 11.6 and 25.1 RLU with *E9-RbcS* and *35S* terminators. In contrast, the *alcSynth-34S promoter* has shown a wide array of expression activity similar to the constitutive promoters; the expression with *HSP* terminator (337.9 RLU) is 7.2 times higher the expression that *LEC2* terminator provides (46.7 RLU) (**Figure 4C**). Both *pOp6-35S* and *lexA-35S* have a basal expression of around 10 RLU (when combined with *HSP*); however, *lexA-35S* has demonstrated ∼11-fold activation amplitude (111.6 RLU), while *pOp6-35S* had only three times activation amplitude (25.1 RLU). Combined with *HSP* terminator, *alcSynth* basal expression was very high (250.3 RLU), leading to only 1.3 times of activation (337.9 RLU) upon treatment with ethanol.

**Figure 4.**
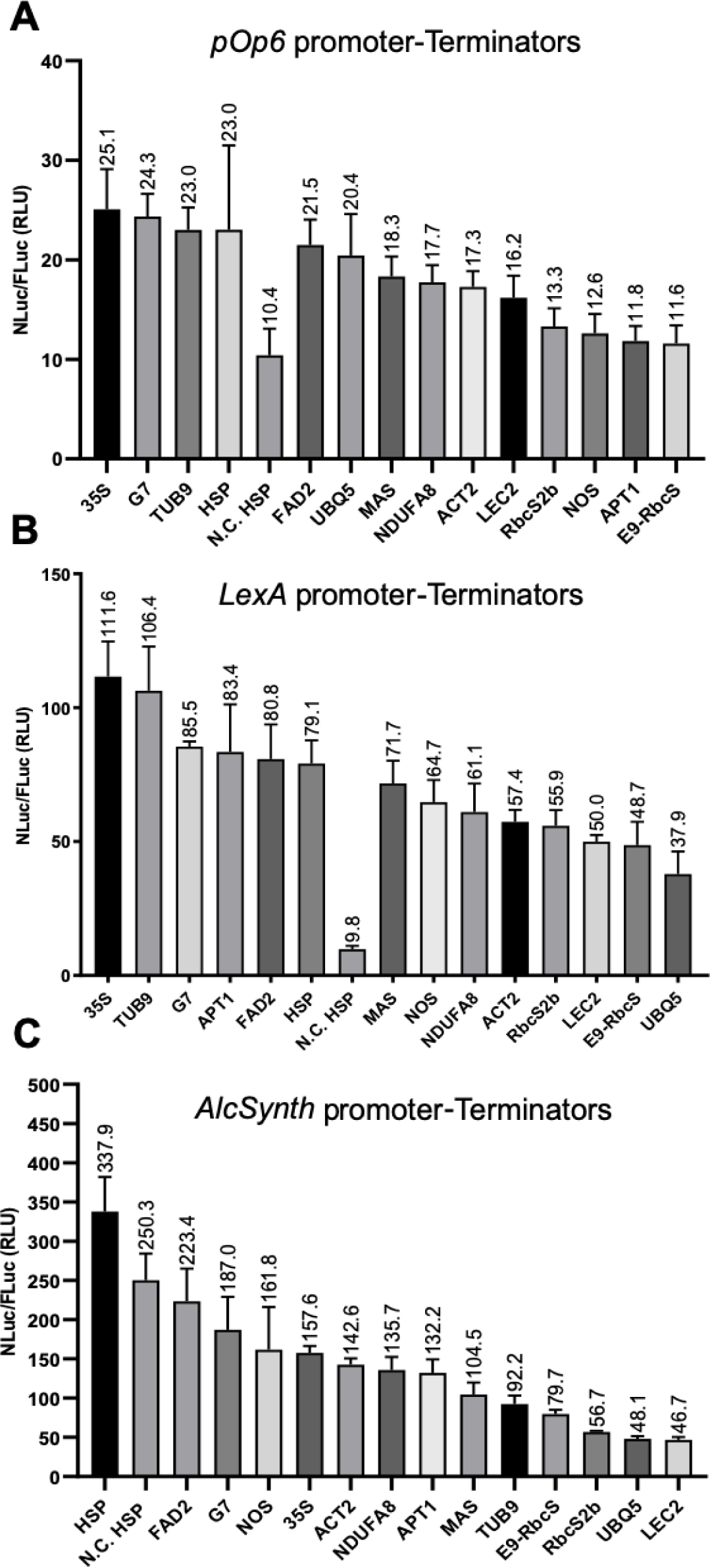
Characterisation of plant inducible promoters with the library of terminators. The characterisation was carried out with PEG-transformation of leaf mesophyll protoplasts from Arabidopsis in 96-well plates. The expression strength was assayed in 96-well plates in a plate reader measuring the Nano luciferase activity normalised by Firefly luciferase. Terminators were characterised combined with the inducible promoters: *pOp6-35S* (**A**), *lexA-35S* (**B**), or *alcSynth-34S* (**C**) inducible promoter. The construct we used was *UBQ10:fluc:UBQ5-UBQ10:Transactivator:HSP-Promoter:nluc:HSP* in pGreen based vector, where ‘Transactivator’ corresponds to *LhGR* (A), *sXVE* (B) and *AlcR* (C) and ‘Promoter’ to *pOp6-35S* (A), *LexA-35S* (B) and *alcSynth-34S* (C). Chemical induction was achieved with 2.5μM DEX for (A), 5μM β-estradiol for (B), or 0.1% EtOH for (C). Nluc = Nano and Fluc = Firefly luciferase activity, bars show normalised luciferase activity values in mean ± SE. N.C. = negative control (uninduced cells).

To gain further insights about how these terminators influence gene expression, we investigated candidate primary sequences that have been implicated in gene regulation via RNA processing or stability or translation efficiency (**Figure 5**). Two strong terminators, *FAD2* and *HSP*, were dissected for putative functional sequences by creating a deletion series.

**Figure 5.**
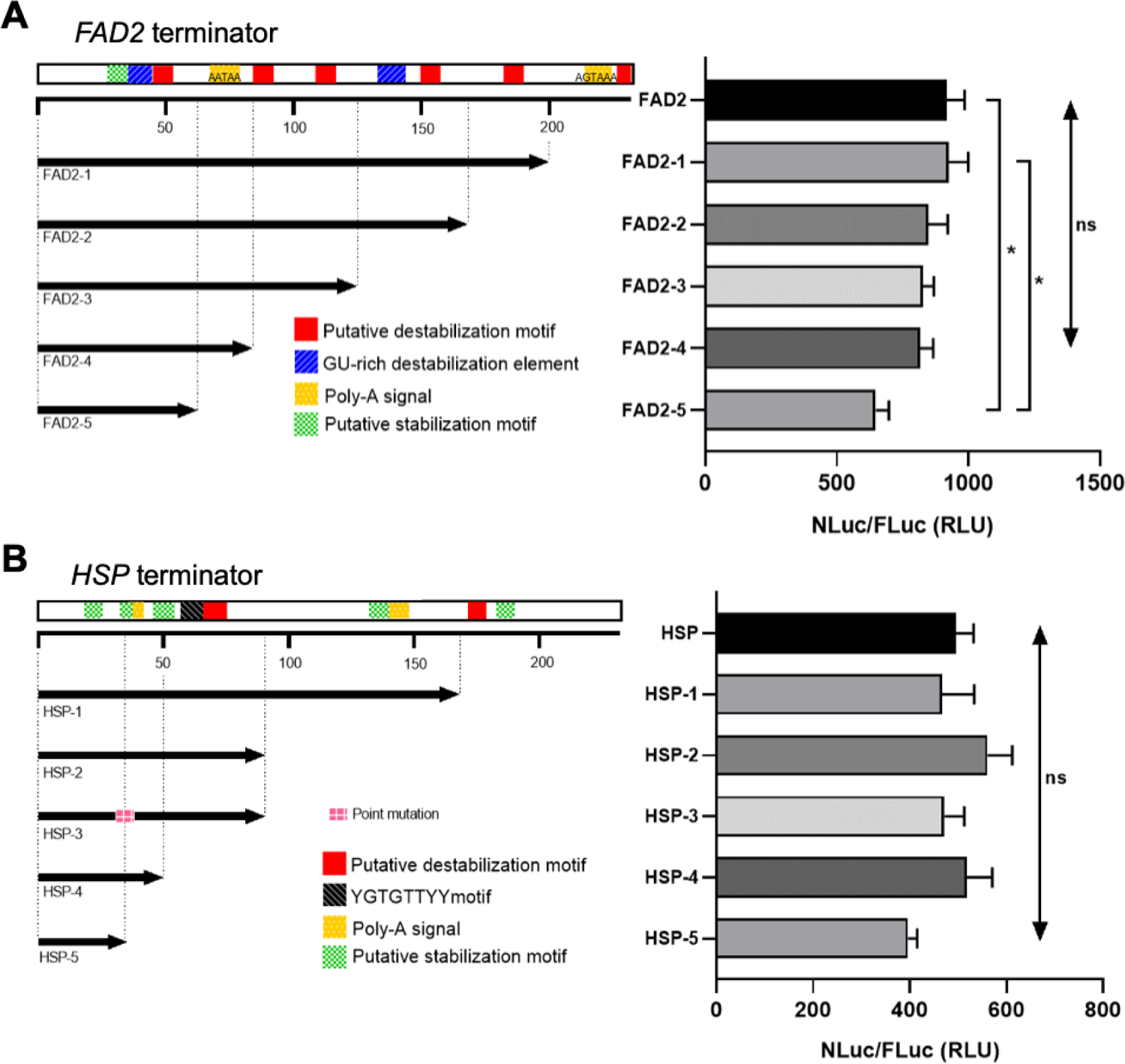
Deletion series of HSP and FAD2 terminators. Five length deletion series were constructed for *HSP* and *FAD2* terminators. The deletions removed putative motifs that could affect their function to influence the gene expression level. Putative Stabilisation/Destabilisation Motifs (Narsai *et al.* 2007), while the rest were identified using RegRNA2.0 and PASPA online tools. Transformation of the construct was performed with PEG-transformation of Arabidopsis leaf mesophyll protoplasts in 96-well plates. The expression activity of the constructs was assayed using a plate reader, measuring the Nano luciferase activity normalised by Firefly luciferase. The construct we used was *UBQ10:nluc:Terminator Part:UBQ10:fluc:UBQ5*, housed in a pMAP vector. Nluc= Nano and Fluc=Firefly luciferase activity, bar graphs show luciferase activity values in mean ± SE. p < 0.05, one-way ANOVA and Tukey’s HSD test.

Putative RNA stabilisation and destabilisation hexamers, Poly-A signals and other regulatory motifs were identified, and we constructed a deletion series to isolate the effects of these motifs. The stabilisation and destabilisation hexamers correlate with mRNA stability (70); the YGTGTTYY motif was found to be necessary for efficient formation of mRNA 3’ termini (71), and Musashi binding element represses the translation of mRNAs in mammalian stem cells or activates translation in maturing Xenopus oocytes (72).

Five different terminator variations were generated for each terminator. For the *FAD2* terminator, Sequence 1 is the full length (200bp) terminator, while Sequence 2 removed the second Poly-A signal and a putative destabilisation signal (PDS) (**Figure 5A**). Sequence 3 resulted from an additional deletion of a PDS and a Musashi binding element (MBE).

Sequence 4 removed another PDS from Sequence 3. Further shortening of the terminator by removing two more PDS gave rise to Sequence 5, and finally, elimination of the first Poly-A resulted in Sequence 6. Concerning the *HSP* terminator, the first dissection removed two overlapping putative stabilisation signals (PSS) and two overlapping PDS (**Figure 5B**). The second deletion of a 73bp segment removed the second Poly-A signal and a PSS to generate the second part. The *HSP* Sequence 3 had a mutation in the Poly-A site converting AATAAA to AAgcAA and removing a segment with a YGTGTTYY motif and two overlappings PDS created part four. Finally, the last part results from the deletion of the Poly-A signal and two PSS. The variation in the terminator activity was evaluated transiently in Arabidopsis mesophyll protoplasts. Statistical analysis of the results showed only a significant difference between *FAD2* Sequence 5, which is the shortest part with no Poly-A site, and the intact *FAD2* as well as FAD2 Sequence 1. Concerning the rest of the *HSP* series as well the FAD2 series, there was no significant difference in the reporter expression level in Arabidopsis protoplasts.

### Promoters and terminators combinatorially regulate gene expression

Comparing two constitutive and three inducible promoters in combination with the whole set of terminators, respectively, we noticed that the gene activation function of a promoter or a terminator is not independent or additive, but they have combinatorial effects. A clear example of such promoter-terminator interaction is seen with the *NOS* terminator, which in combination with *UBQ10* results in weak, *NDUFA8* strong, *lexA* medium, *pOp6* weak and *alcSynth* medium expression (**Figure 6**). From the 14 terminators characterised, only four exhibited stable behaviour: *FAD2* and *HSP* were consistently on the strong side, while *LEC2* and *RbcS2b* tending towards the weak side. The terminators paired with the inducible promoters *pOp6-35S* and *lexA-35S* resulted in negligible interaction as they follow a specific pattern (strong/medium/weak), with the exemption of *UBQ5*, *APT1* and *NOS*. On the other hand, *AlcSynth-34S* does not follow this pattern.

**Figure 6.**
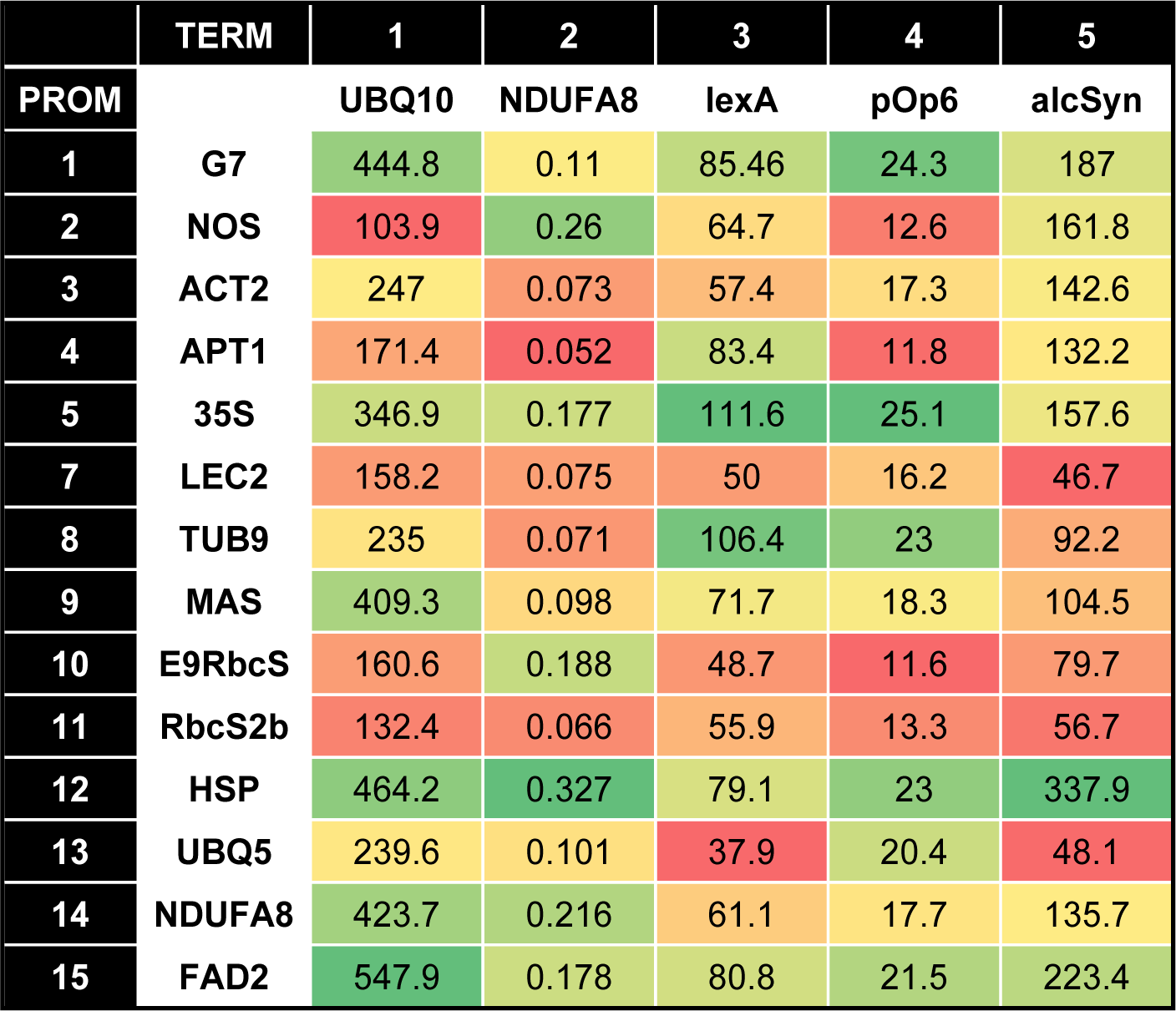
Promoter-terminator interactions in gene regulation. Heat map showing how the combination of different promoters and terminators affects the expression level of nanoluciferase normalised by firefly luciferase. The green colour indicates high activity, while red low. The colour scale is set independently in each promoter series.

## DISCUSSION

### MAPS: A new synthetic biology framework to engineer plant systems

In the present work, we established the Mobius Assembly for Plant Systems (MAPS), which is an adaptation of the Mobius Assembly (9) for plant gene expression. MAPS is a fast and user-friendly DNA assembly platform based on small plant binary vectors, and it comes with a characterised part toolkit for plant expression. We have introduced a new feature for Mobius Assembly to facilitate combinatorial DNA library assemblies. This new feature – MethyAble – exploits *in vitro* DNA methylation to directly feed Phytobricks in the Level 2 constructs.

For plant genetic engineering, small binary vectors are vital in enhancing efficiency in cloning and transformation. Initially, we have used the pGreen backbone for protoplast transformations; however, they had transformation issues in *E. coli* TOP10 cells with large constructs and proved inefficient for transforming plant cell cultures. As we wanted a reliable core vector in our kit for both cloning and transformation applications, we developed a new binary vector pMAP. pMAP bears a fused replication of origin of pUC with pWKS1 from a *Paracoccus pantotrophus* cryptic plasmid, and it is suitable for experiments that demand both high DNA yield (e.g. cloning and protoplast transformation) and transformation efficiency (e.g. cell cultures). This vector was highly efficient for cell culture transformations and was used to characterise dissected terminator parts in both Arabidopsis protoplasts.

To test the functionality of the MAPS vectors, we used them in combination with five commonly used Agrobacteria strains to transform the Arabidopsis MM1 cell line. We showed that the vector selection is of crucial importance for the cell culture transformation, with pLX based and pMAP vectors outperforming pGreen vectors. Among pLX and pMAP, there was no considerable difference in the strains C58C1*, LBA4404 and AGL1, while pMAP demonstrated better activity in GV3101. Agrobacterium chromosomal background played a vital role in the infection of the MM1 Arabidopsis cell line. LBA4404 (Ach5 background) outperformed all the strains with C58 background (C58C1*, AGL1 and GV3101). On the contrary, in Arabidopsis floral dip transformation, GV3101 was shown to have higher transformation frequencies than LBA4404 (73), while in transient assays, C58C1* found to be the best strain either for leaves (74) or seedlings (75). Among the strains with C58 background, AGL1 provided the lowest infection showing that the disarmed virulent vector is another factor that affects the transformation efficiency. The other two C58 strains (GV3101 and C58C1*) exhibited strong transformation efficiency, with GV3101 being slightly more efficient when harbouring the pMAP vectors. Factors that can explain the results are the copy number of the vector, the segregational plasmid stability, and the T-DNA borders. pLX and pGreen vectors have a medium and low copy number origin of replication for Agrobacteria, respectively. pMAP, on the other hand, resulted in plasmid concentrations between pLX and pGreen. The higher the copy number, the more copies of a construct can be transferred to the cells. pGreen’s origin of replication, pSa, does not include the stability regions necessary for maintaining the plasmid (50, 100). On the other hand, pLX and pMAP origins of replication (BBR1 and WKS1) were derived from cryptic plasmids that are stable without stability regions (76, 101). As there is no selective pressure during Agrobacterium transformation, possibly, pGreen plasmids are not 100% maintained in the agrobacteria strains. Also, even though WKS1 is thought to replicate at a lower copy number than BBR1, pMAP exhibited similar or stronger transformation efficiency than pLX, which might be attributed to its better segregational plasmid stability. A third factor that can affect the transfection is the T-DNA borders of the binary vector. pGreen has minimal synthetic LB and RB sequences, derived from pTiT37 and an Overdrive from the LBA4417 plasmid (50), while pLX and pMAP have extended LB and RD T-DNA borders, including the overdrive from the plasmid pTiA6 (54).

### New libraries of short promoter and terminator parts

Furthermore, we generated and characterised a collection of short promoter (17 constitutive and three inducible) and 14 terminator standard parts for plant expression. Ten of the promoters and six of the terminators were newly isolated in the present study, and they have a sequence length between 300-500bp (promoters) and 200bp (terminators). Until now, little attention has been paid to the size of the developed plant parts. Available in repositories (e.g. MoClo plants, GB plants and GreenGate) are promoters and terminators up to 4kb. The large size of the expression elements can burden the multigene delivery in plants as the large size of the binary vectors will lead to plasmid instabilities and low transformation efficiency or incomplete/truncated transformation (38, 45). There are indications that short plant terminators are adequate for transcription termination. Notably, it was shown that transcripts with 3′ UTRs longer than 300 bp do not improve mRNA stability (70). In another study, ten randomly selected mRNA sequences from Arabidopsis showed an average 3’UTR length of 209bp (76). Our short terminators were found to have matching or surpassing expression activity than the commonly used terminators (e.g. *NOS*, *35S*). Additionally, there are already examples of short, strong constitutive promoters from Arabidopsis, such as *AtTCTP* (0.3 bp (77)) and *AtSCPL30* (0.45bp (78)). A recent study was also introduced short synthetic promoters for plant expression (79).

With this study, the plant toolbox is further equipped with short promoters with a range of expression levels, desirable for different applications such as overexpression of transgenes or expression of resistance cassettes. An *Actin 2* promoter (787bp) was characterised in MoClo and GB2.0 toolboxes, showing a comparable activity to the NOS promoter (80, 81). The shorter version we made in this study (340bp) exhibited similar expression activity (**Figure 3**). More specifically, in combination with the HSP terminator, the *NOS* and *ACT2* promoters had 1.4 and 4.6 RLU, respectively, while the corresponding expression from the GB2.0 versions was 2.61 and 4.36 RLU (40). *UBQ11* and *UBQ4* promoters showed strong expression, and they performed better than the commonly used *35S* and *OCS*. Sequence analysis through the online programs PLACE and PlantCARE showed that the *UBQ11* promoter contains 11 CAAT-box and 6 DOFCOREZM, while the *UBQ4* promoter harbours 9 CAAT-box and 7 DOFCOREZM elements (**Supplementary Figure S2**). DOFCOREZM elements are the binding sites of Dof1 and Dof2 TFs, which were found to enhance gene expression (82). CCAT-box also influences expression efficiency (83). The *UBQ10* promoter is a well-known and commonly used constitutive promoter, and it exhibited the highest expression in our protoplast assay, ∼20 and ∼11 times higher than the 35S promoter, combined with *NDUFA8* and *HSP* terminators, respectively. On the contrary, in the part characterisation in *Nicotiana benthamiana* leaves, the *35S* promoter had the highest activity as shown by the MoClo and GB2.0 publications (80, 81). Similarly, the *OCS* promoter had strong activity in our study but low activity in the characterisation in tobacco leaves (80). Consequently, some genetic parts behave differently in different chassis, and we should not generalise characterisation results among different plant species or expression systems (whole plants, transient leaf expression systems, protoplasts, or cell cultures). Even though the *ACT7*, *TUB2*, *TUB9*, *APT1*, *ACT2* and *LEC2* promoters had lower expression activity, they performed better than *NOS*, the commonly used promoter to drive the antibiotic/herbicide resistance for transgene selection. Therefore, they can be used to express selection markers or any experiments that do not require high expression levels of the transgenes.

### Terminator affects gene expression

Comprehensive characterisation of the terminators using the high throughput protoplast transient assay that we developed underscored the strong influence of the terminators on gene expression. It is a common practice to rely on promoters to control gene expression level and neglect the influence of terminators. For example, to compensate the loss of expression after the domestication of the synthetic plant promoter *G10-90* promoter, researchers replaced it with the *AtRPL37aC* promoter (84). However, they were using the *psE9-RbcS* terminator, which we found to drive low gene expression; the gene expression could have been restored with a more activating terminator instead. Terminators could alter the gene expression 5.3, 6.3 and 7.3 times when combined with *UBQ10*, *NDUFA8* and *alcSynth* promoters.

Terminators can alter gene expression in multiple ways. 3’UTR participates in quality control in the post-transcriptional regulatory mechanism for eukaryotic genes through Nonsense-mediated mRNA decay (57, 58). Genome-wide analysis of the transcript degradation profiles in Arabidopsis revealed specific sequence motifs at 3’ UTR that stabilise or destabilise mRNA (59). Several motifs in the 3’UTR might be involved in polyadenylation or in mRNA decay (60). Strong gene expression is linked with short poly(A) tails, while poly(A)-binding proteins can facilitate both the protection and degradation of mRNA (61). In Arabidopsis, the poly(A) tail was found to block RNA-DEPENDENT RNA POLYMERASE 6 (RDR6) from converting aberrant mRNAs into substrates for degradation (62). Unpolyadenylated transcripts derived from terminator-less constructs or readthrough mRNAs from transgenes with strong promoters are subjected to (RDR6) mediated silencing (63).

To gain insights into how the terminators influence gene expression, we dissected different domains of the *HSP* and *FAD2* terminators based on sequence elements and known RNA processing or motifs affecting translation efficiency. We mapped putative stabilisation and destabilisation hexamers, Poly-A signals and other regulatory motifs known on the terminators and constructed a deletion series to isolate the effects of these motifs (**Figure 5**). Only removing the *FAD2* Poly-A site was able to considerably reduce the expression in Arabidopsis protoplasts, revealing that it is a key driver of this terminator activity. For the other tested motifs, no conclusive correlations to the expression activity were made, as there were no statistically significant effects on the readout of the luciferase activity. These results suggest that the known sequence motifs in terminators affecting gene activity cannot fully explain the strong influence of terminator cast in gene regulation. This was further supported by another observation we made that the relative strength of a promoter was dependent on the terminators it was paired with.

Chemically inducible systems are popular tools to control the timing and amplitude of transgene expression. β-estradiol, Dex, and ethanol-inducible systems were developed almost 20 years ago and have been used in many studies since then (69). However, no work has been reported about enhancing the target gene expression using different terminators. In this study, we attempted to modulate the gene expression ranges of the inducible promoters by exchanging the terminators. We found that *lexA* has a 2.9 times difference in the expression level when combined with different terminators, while *pOp6* showed 2.2 times difference (**Figure 4**). In contrast, the *alcSynth* led 7.2 times difference in reporter gene expression depending on the terminators. *pOp6* and *lexA* are fully synthetic and comprised of a minimal promoter and a tandem repeat of the specific transcription factor binding sites; in contrast, the alcohol inducible system (36) drives its promoter from the natural promoter found in *Aspergillus nidulans*. We therefore propose that *alcSynth* promoter still harboured sequences that mediate interaction with the terminators (**Supplementary Figure S3**), whereas *pOp6* and *lexA* do not.

The Dex and Estradiol systems had similar basal expression levels of ∼10 RLU (with the *HSP* terminator) in the protoplast system and with our hands; however, *lexA-35S* has demonstrated ∼10-fold activation while *pOp6-35S* only 3-fold (**Figure 4A and B**). The *alcSynth* promoter, on the other hand, had a high basal expression (250.3 RLU with the *HSP* terminator) with 1.3 times activation (**Figure 4C**). A possible explanation for the leakiness of the ethanol system might be TMV Ω; the addition of the TMV Ω in the *LhGR-pOp6* system exhibited slightly increased basal expression (85). Also, as the whole *alcA* promoter is directly fused to the minimal *35S* or *34S* promoter, constitutive transcription elements present in the sequence might be responsible for the leakiness. Synthetic design of the ethanol inducible promoter, in which only *alcR* binding sites are present separated by random spacer sequences and fused to a minimal plant promoter, could reduce the leakiness and improve the stringency of the gene regulation. Taken together, the estradiol and Dex inducible systems worked well in our protoplast systems, and more inducible systems, such as optogenetics or other chemical inducible systems, are anticipated to enrich the synthetic biology toolkit to build circuitry gene expression control in plant systems.

### Promoter-terminator interactions in gene regulation

We have observed clear interactions between promoters and terminators, as the terminators drove relatively different expression levels when combined with different promoters and *vice versa* (Figure 6). The *NOS* terminator is a case in point; It behaves as a weak terminator in conjunction with the *UBQ10* and *lexA* promoters, yet as a strong terminator with the *NDUFA8* promoter and medium terminator with the *alcSynth* and *pOp6* promoters. Only four out of the 14 *terminators* exhibited stable behaviour: *FAD2* and *HSP* were always strong while *LEC2* and *RbcS2b* were always weak. Therefore, a terminator can have a different impact on different promoters. Terminators performed differently combined with the inducible promoters *pOp6-35S* and *lexA-35S* than the rest of the promoters. These combinations resulted in a more uniform expression range and exhibiting similar terminators strengths (low/high). *LexA-35S* and *pOp6-35S* inducible promoters are synthetic and comprise only the minimal *35S* promoter and bacterial DNA binding sites. Hence they do not include other regulatory elements that can interact with the promoters and further manipulate gene expression. In contrast, *alcSynth* includes the intact *alcA* promoter, consequently behaves similarly to the other promoters and contains more regulatory elements.

The synergistic regulation between natural promoters and terminators can be explained by the direct interaction of the terminator with the promoter (RNA or chromatin looping), which can further influence gene expression. Gene looping is the physical connection of the terminator with the promoter region of a gene (86), mediated by nucleic acid-binding proteins (87). It can influence gene expression through transcriptional memory (88), intron-mediated modulation of transcription (89), transcription directionality (90), reinitiation of transcription (91), and transcription termination (92, 93). Moreover, it was recently shown that the absence of a terminator caused higher DNA methylation on the promoter region and reduced the transgene expression compared to constructs with a terminator, indicating an alternative role in transcriptional gene silencing (94).

It was shown previously that double terminators increase expression (95, 96). Building on the double terminator observations, Diamo and Mason showed a synergistic enhancement in the double terminator combinations (97). The effect of the double terminators was partially attributed to a reduction of RDR6-mediated silencing. mRNA read through from “leaky” terminators under the influence of a strong promoter triggers RDR6-mediated silencing.

However, this explanation cannot sufficiently explain the synergistic effects the terminators cast, especially combined with weak promoters or double terminators. Meanwhile, interaction of the terminators with the promoters and the transcription initiation machinery could. More in-depth work, such as RNA/chromatin structure assay and *a priori* dissection of the promoter and terminator sequences in influencing gene activity, is required to verify the theory of looping or any other mechanisms on the roles of plant terminators on transcriptional or translational regulation.

## Perspectives/conclusion

Plant synthetic biology lacks characterised parts to develop more predictable biosynthetic pathways and high-throughput methods to perform systematic genetic part characterisation. There is also a need for characterised genetic elements in different plant chassis, for example, Arabidopsis, tobacco, and barley. In this work, we equipped the plant community with a DNA Assembly (MAPS), new compact binary vectors for general plant expression and a library of short characterised genetic elements for Arabidopsis. Throughout this work, we underlined the power of terminators in controlling gene expression, which is neglected, and attention primarily focuses on promoters. We also showed that terminators act in synergy with the promoters, and it is incorrect to assume that a promoter or a terminator can always be strong or weak. Further studies need to be conducted to elucidate the mechanisms of these interactions.

In the plant community, there is also the tendency to use the same *E. coli* and Agrobacterium strains, binary vectors, cloning and transformation methods, for all experiments conducted within a lab, assuming that their efficiency is independent from each other. Here we showed that the combination determines the overall efficiency of an experiment in non-additive manners. The successful cloning of complex multigene constructs, plant transformation and expression is affected by the bacterial stains and the binary vectors. Cell strains (*E. coli* and Agrobacteria) can be responsible for low transformation efficiency and plasmid instabilities. Simultaneously, binary vectors can also contribute to plasmid instabilities, low transformation efficacy and protein expression. Consequently, all of these factors need to be considered in combinations when working with complex constructs in plant engineering.

## SUPPLEMENTARY MATERIALS

### Results

#### Plasmid structural stability

Structural stability of the plasmids was evaluated during propagation in *E. coli* and Agrobacterial strains. The 3-TU insert *UBQ10:LhGR:HSP-pop6:nluc:HSP-UBQ10:fluc:UBQ5* was cloned either in pGreen, pLX, pLXBBR1mut or pMAP-based Mobius Assembly Vectors. pLXBBR1mut was an attempt to increase the copy number of pLX vectors for applications that demand a high amount of DNA. pGreen based vectors exhibited good stability in DH10B (5/5), Dh5α (5/5), and TOP10 (4/5) but poor stability in JM109 (0/5) and NEB stable (0/5) strains according to the restriction digestion profiles and sequencing results **(Supplementary Figure S1**).

**Supplementary Figure S1.**
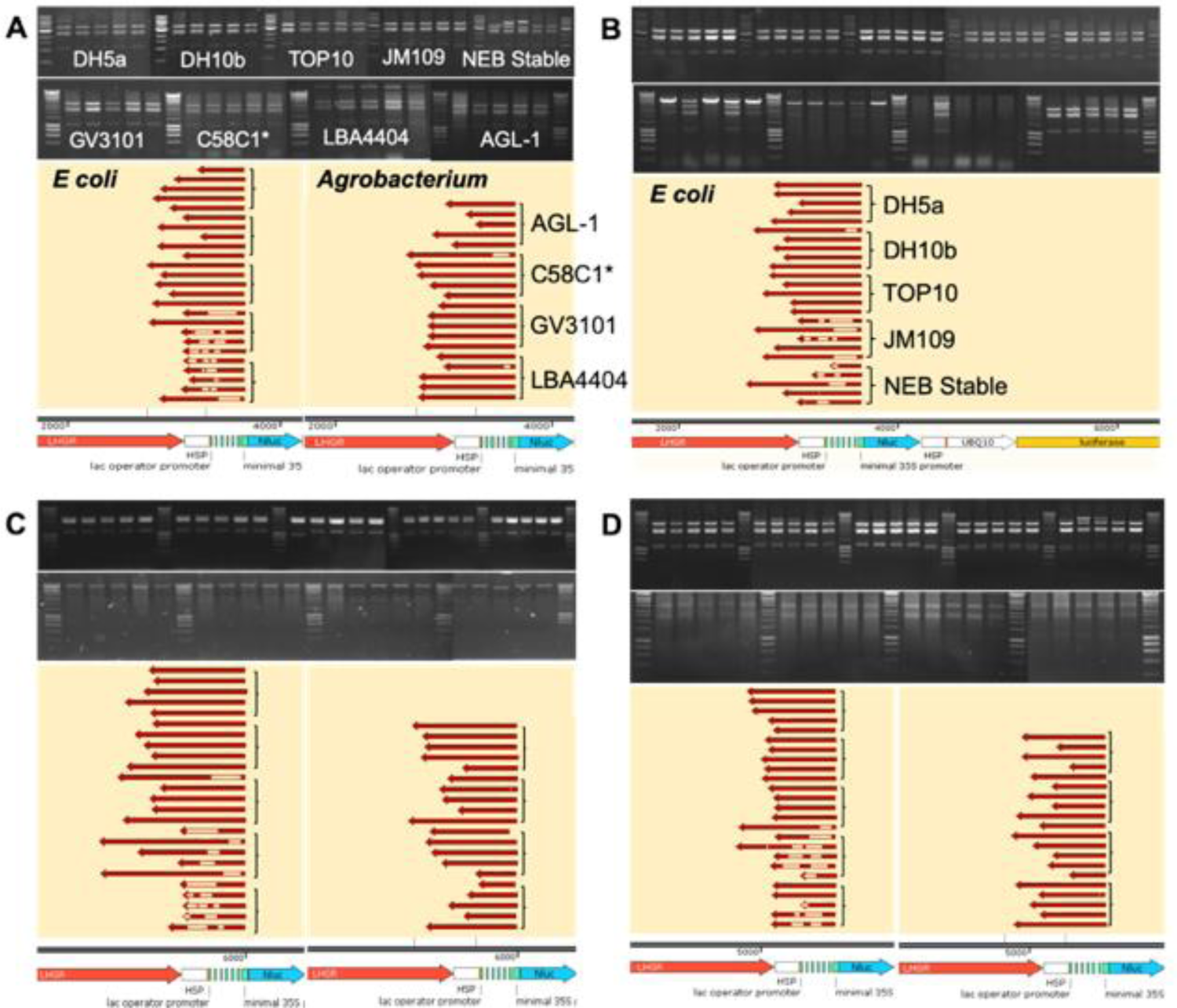
Plasmid structural stability analysis. Stability was tested by electrophoresis in 1% agarose gel after digestion with *PstI*-HF and then by Sanger sequencing of the tandem short repeats of the pOp6 sequence. The stability of the construct *UBQ10:LhGR:HSP-pOp6:nluc:HSP-UBQ10:fluc:UBQ5* was evaluated in both *E. coli* (DH5α, DH10B, TOP10, JM109, and NEB stable) and *A. tumefacience* strains (AGL-1, C58C1*, GV3101, and LBA4404) housed by the binary vectors: pLX (**A**), pLX-BBR1mut (**B**), pGreen (**C**) and pMAP (**D**). Alignment of the sequencing data was performed with Snapgene. The concentration of pGreen generated in Agrobacteria is low; consequently, we used high volume of plasmid DNA, but the digestion was incomplete. Low concertation of pMAP also resulted in low visibility of the second band.

Instabilities were detected as random deletions in the pOp6 repetitive sequence. pLX based plasmids found stable in DH5α (5/5), DH10B (5/5) and TOP10 (5/5) strains; however, they also showed poor stability in NEB stable (0/0), and JM109 (0/0) strains with deletions in the pOp6 domain. A similar pattern was observed for the pLX BBR1mut version for *E. coli strains*, as they were stable in DH5α (5/5), DH10B (4/5) and TOP10 (5/5) strains, yet unstable in JM109 (1/4) and NEB stable (1/4) strains. Lastly, the pMAP vector was stable in DH5α (5/5), DH10B (5/5) and TOP10 (5/5) and also unstable in NEB stable (2/5) and JM109 (0/5). Regarding the Agrobacteria strains AGL1 (4/5), GV3101 (5/5), LBA4404 (5/5) and C58C1* (5/5), no major instability problems were detected in the pGreen vector. pLX was also stable in Agrobacteria with AGL1 (5/5), GV3101 (5/5), LBA4404 (4/5) and C58C1* (4/5). pMAP was stable in Agrobacteria as well, having all the colonies correct (AGL1 (5/5), GV3101 (5/5), LBA4404 (5/5) and C58C1* (5/5). Nevertheless, only the AGL1 strain could propagate stable plasmids for the pLXBBR1mut version, which has a higher plasmid copy number. Severe instability issues were observed in GV3101, LBA4404 and C58C1* strains with aberrant restriction digestion patterns and even plasmid loss, rendering the high copy BBR1mut plasmids unsuitable for propagation in Agrobacteria.

### Methods

#### Structural plasmid stability

The construct *UBQ10:fluc:UBQ5-UBQ10:LhGR:HSP-pop6:nluc:HSP* was used for the stability studies and cloned either in pGreen based, pLX BBR1 based, pLX BBR1mut or pMAP based Mobius Assembly vectors and transformed in DH5α, DH10B, TOP10, JM109 or NEB stable *E. coli* strains. LB cultures, after 24h of incubation, were used for plasmid isolation. Structural plasmid stability was tested first electrophoretically in 1% agarose gel after digestion with PstI-HF (NEB) and then by Sanger sequencing of the tandem short repeats of the pOp6 sequence. Next, the Agrobacteria strains AGL-1, C58C1*, GV3101, LBA4404 were transformed with a construct from each plasmid with the correct sequence, and the same process was followed to determine the structural plasmid stability.

For pLX-BBR1mut construction, primers were designed to introduce a point mutation (C299 to T299) into the Rep gene of pBBR1 origin of replication to increase the copy number (60). pLX Level 1A and Level 2A plasmids were PCR amplified into two parts using the primers carrying the mutation and the Assembly Linker 1 primers. Gibson assembly was employed to reconstruct the plasmids, and after transformation, colonies with bright pink and yellow colour were, selected for the pLX BBR1mut version.

### Discussion

#### The effect of vectors and bacterial strains on plasmid stability

Our results indicate that structural plasmid stability is an essential factor to consider when working with constructs susceptible to instabilities. The insert employed to test structural plasmid stability (UBQ10:LhGR:HSP-pOp6:nluc:HSP-UBQ10:fluc:UBQ5) is prone to instabilities since it is large (∼6.7 kb) and has two direct repeats (two UBQ10 promoters and two HSP terminators). Additionally, the pOp6 promoter consists of six inverted repeats of the lac operator and five inverted spacer repeats. Most of the observed instabilities were deletions in the pOp6 sequence. Structural plasmid analysis revealed that the stability was strain-dependent; The constructs housed by pGreen, pLX, and pMAP vectors were generally stable in DH10B, DH5a, TOP10 strains but unstable in JM109, and NEB stable strains.

Interestingly, even though all the *E. coli* strains we used are recA1^-^ to minimise recombination, they do not perform the same in plasmid stability. Similar conclusions were drawn by Moore *et al*. when they switched to JM109 from DH10Β to deal with instability issues in the engineering of the violacein pathway (1). All the vectors mentioned above were generally stable in the Agrobacteria strains (AGL1, GV3101, LBA4404, C58C1); however, the domains vulnerable to instabilities should be sequenced before any application.

Increasing the copy number of pLX based vectors by mutating the BBR1 origin of replication maintained the stability of the vectors in the *E. coli* strains, which are recA^-^, but abolished it in Agrobacteria strains, except AGL1 (**Supplementary Figure S1B**). AGL1 is recA^-^ strain, which stabilises the recombinant plasmids, hence justifying the results (2). Additionally, all the Agrobacteria strains had a slower growth rate on agar plates (3-4 instead of 2-3 days). Even though BBR1mut was tested before in the C58 Agrobacterium strain to produce β-carotene (3), we do not consider a suitable origin of replication for binary vectors because it could not stably maintain large and complex constructs in commonly used Agrobacteria strains. Overall, a selection of an origin of replication with a bacterium strain that favour plasmid stability should be considered before transforming a construct into plants or plant cells,

**Supplementary Figure S2.**
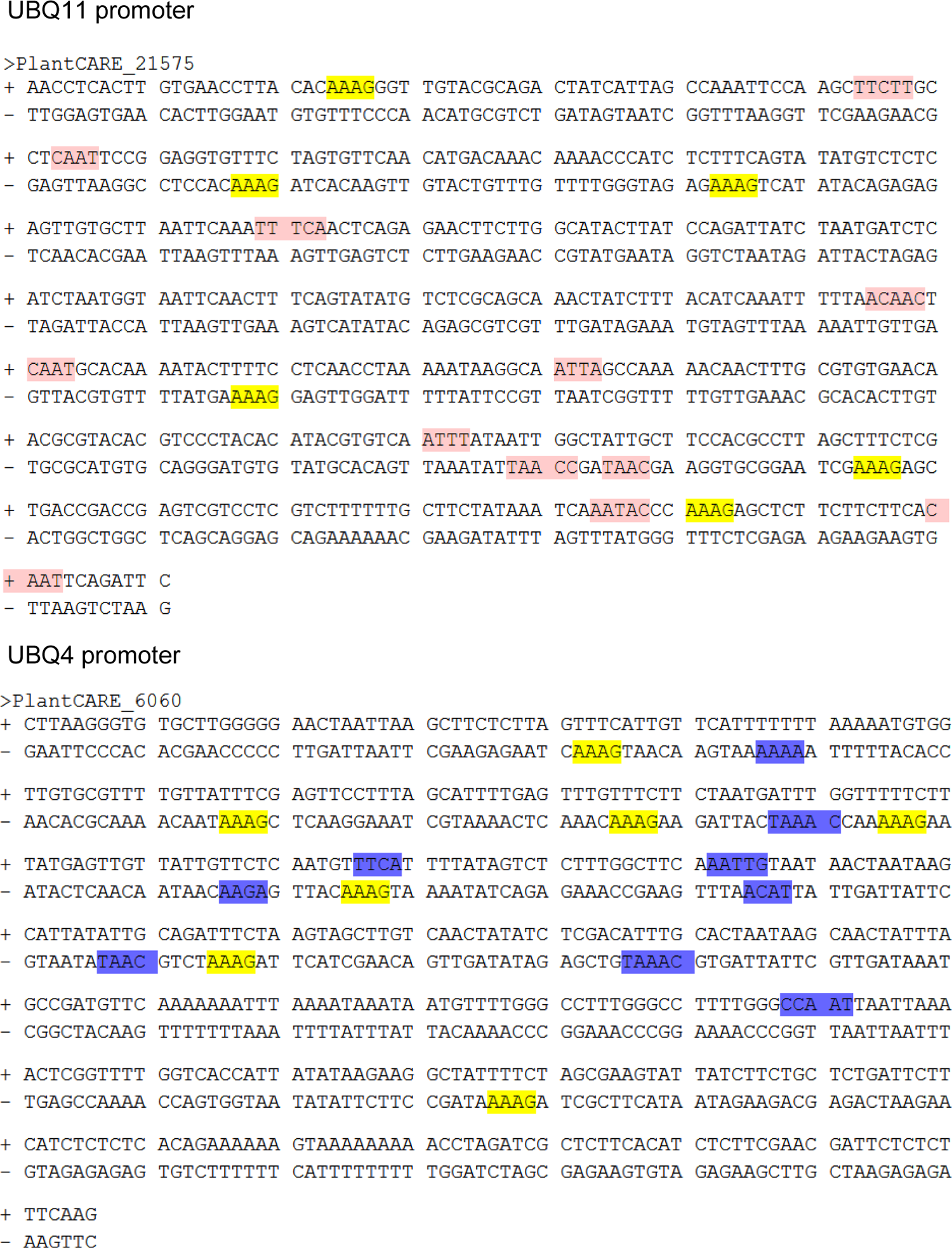
Potential sequence elements in the UBQ11 and UBQ4 promoters linked to increased gene expression. Sequence analysis performed with the online programs PLACE and PlantCARE. Yellow highlighted sequences show the DOFCOREZM elements, pink the CAAT-boxes in the UBQ11 promoter and blue the CAAT-boxes in the UBQ4 promoter.

**Supplementary Figure S3.**
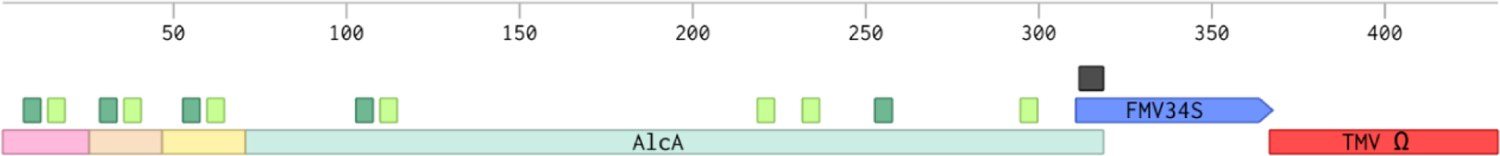
Regulatory elements in the *alcSynth* ethanol inducible promoter. It is a fusion of the alcohol dehydrogenase promoter sequence (alcA) and elements of the alcM (pink), alcR (Orange) and aldA (yellow) with the minimal Figwort Mosaic Virus (FMV) 34S and the translational enhancer from the tobacco mosaic virus 5’-leader sequence (TMV Ω). Deep green is 5’ ➔3’ and pale green 3’ ➔5’ AlcR binding sites. Black box is TATA box.

### Supplementary Tables

**Supplementary Table S1.**
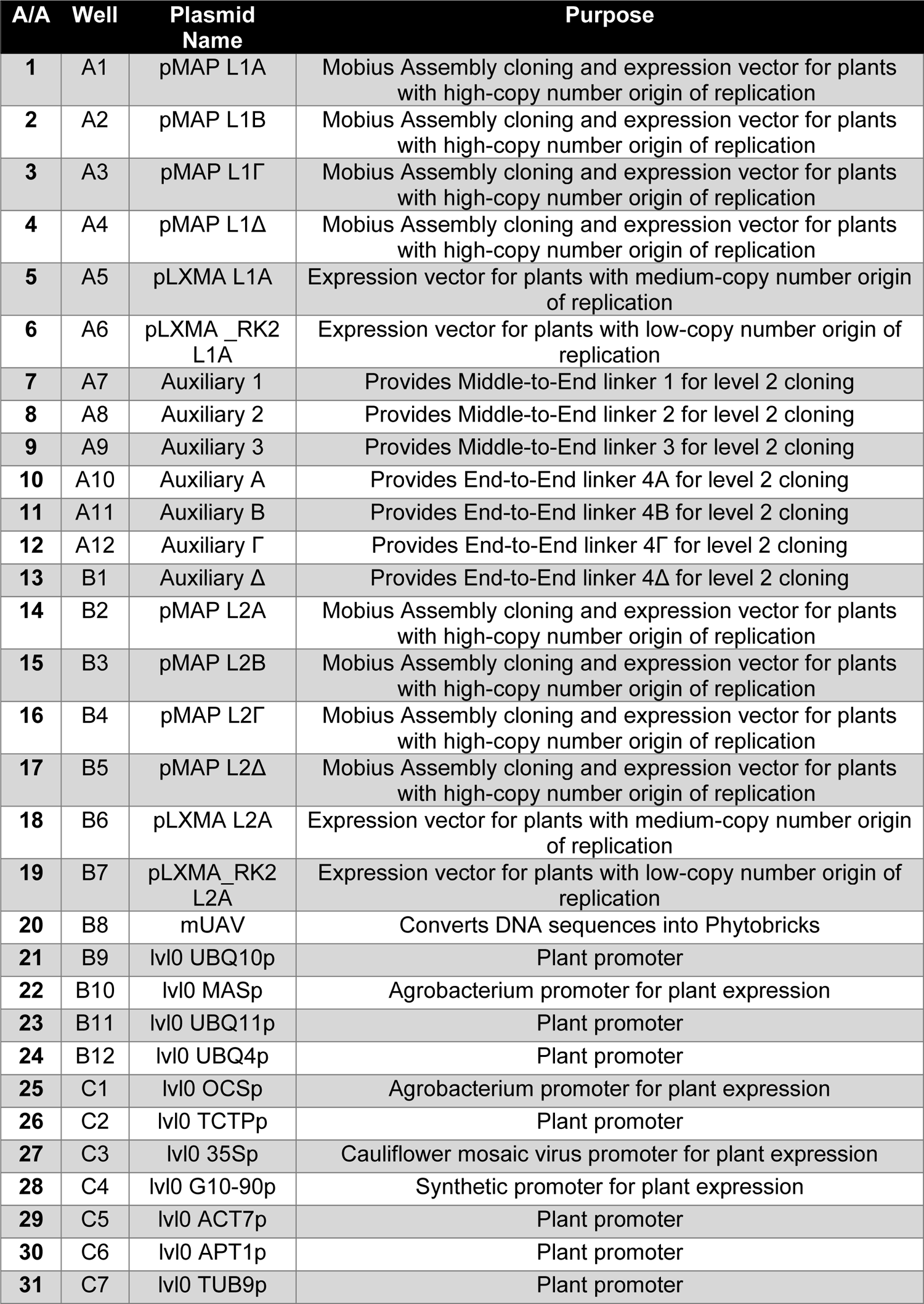
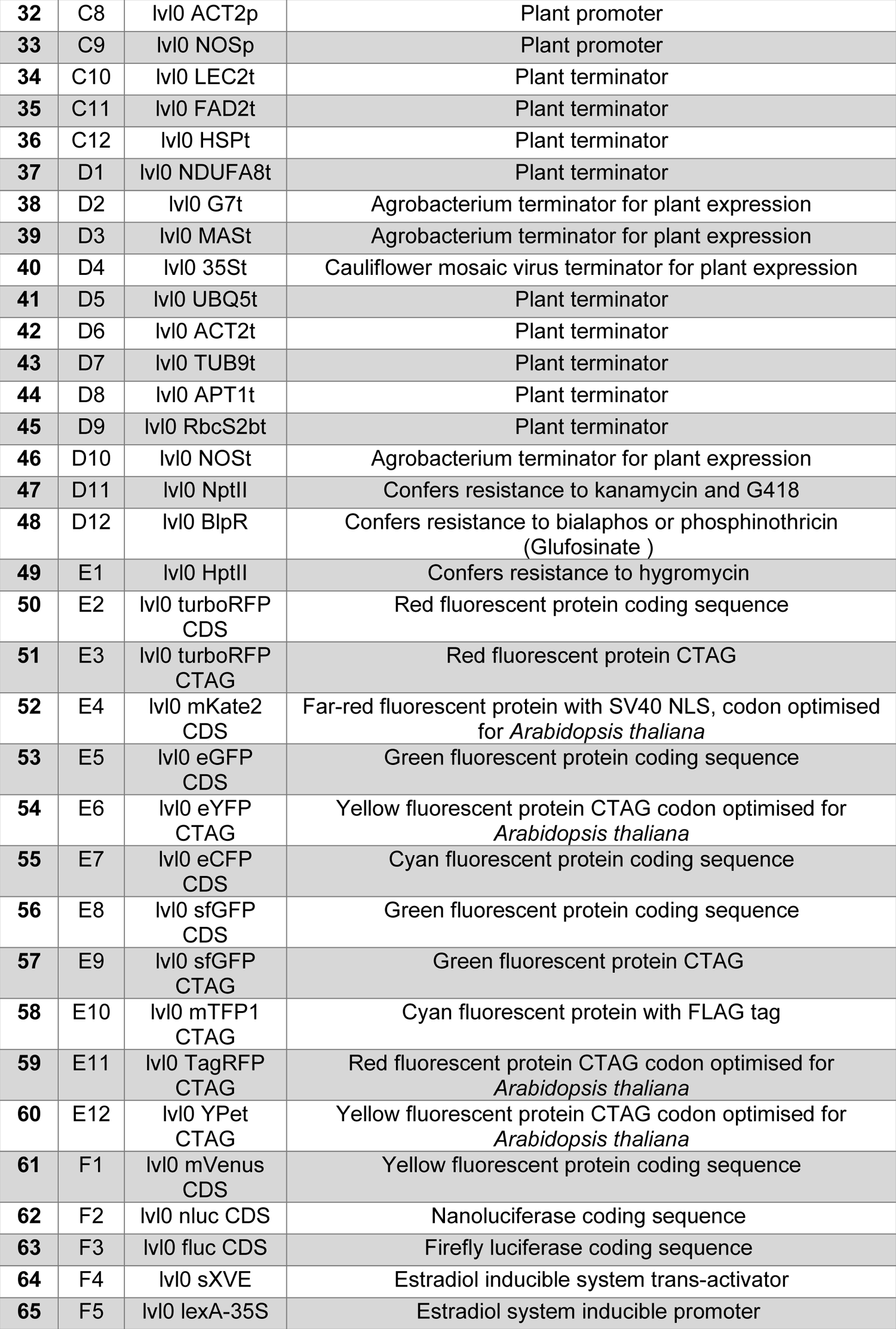
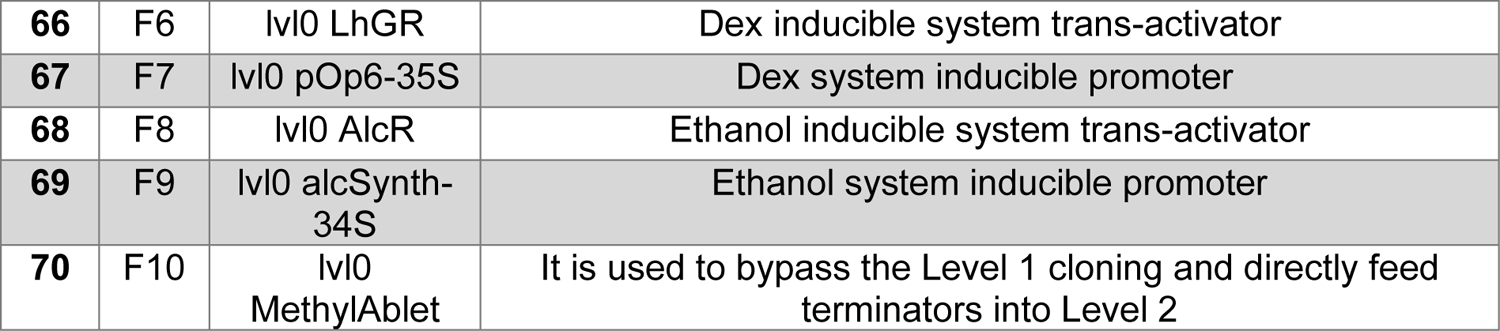
Plasmids contained in the MAPS toolkit. They are available from AddGene (https://www.addgene.org/browse/article/28211394/).

**Supplementary Table S2.** Agrobacterium strains commonly used in plant biotechnology. They are derived from two wild isolates, C58 and Ach5 (4), and they differ in their Ti plasmids. C58 contains two plasmids, the nopaline type pTiC58 and a cryptic one pAtC58, which assists the virulence (5). Temperature-induced loss of pTiC58 plasmid generated C58C1 strain (6). The introduction of the disarmed pTiC58 plasmid (pMP90) led to the generation of GV3101:pMP90 (7). AGL1 also has the C58C1 background and has been engineered to carry a supervirulent, disarmed succinoamopine-type plasmid, pTiBo542, from the Agrobacterium strain A281 (2). C58C1(pTiB6S3ΔT)H derives again from the C58C1 strain and contains a disarmed octopine-type Ti plasmid pTiB6S3 (8) and harbours the pCH32 plasmid, which overexpresses the virulence genes virE and virG (9). LBA4404 has Ach5 as its chromosomal background and contains a disarmed octopine-type plasmid pTiAch5, named pAL4404 (10).

| ***Agrobacterium strain*** | ***Parental strain*** | ***Virulent vector*** | ***Antibiotic Resistance*** | ***Antibiotic Concentration*** |
| --- | --- | --- | --- | --- |
| *AGL1* | *C58* | pAtC58<br>disarmed<br>pTiBo542 | *Rifampicin*<br>*Carbenicillin* | *50 µg/ml*<br>*100 µg/ml* |
| *GV3101* | *C58* | pAtC58<br>pMP90(disarmed<br>pTiC58) | *Rifampicin*<br>*Gentamycin* | *50 µg/ml*<br>*25 µg/ml* |
| *LBA4404* | *Ach5* | pTiAch5, named<br>pAL4404 | *Rifampicin* | *50 µg/ml* |
| *C58C1\**<br>*C58C1(pTiB6S3ΔT)<sup>H</sup>* | *C58* | pTiB6S3<br>pAtC58<br>pCH32 | *Rifampicin*<br>*Tetracycline*<br>*Carbenicillin* | *50 µg/ml*<br>*15 µg/ml*<br>*100 µg/ml* |

**Supplementary Table S3.**
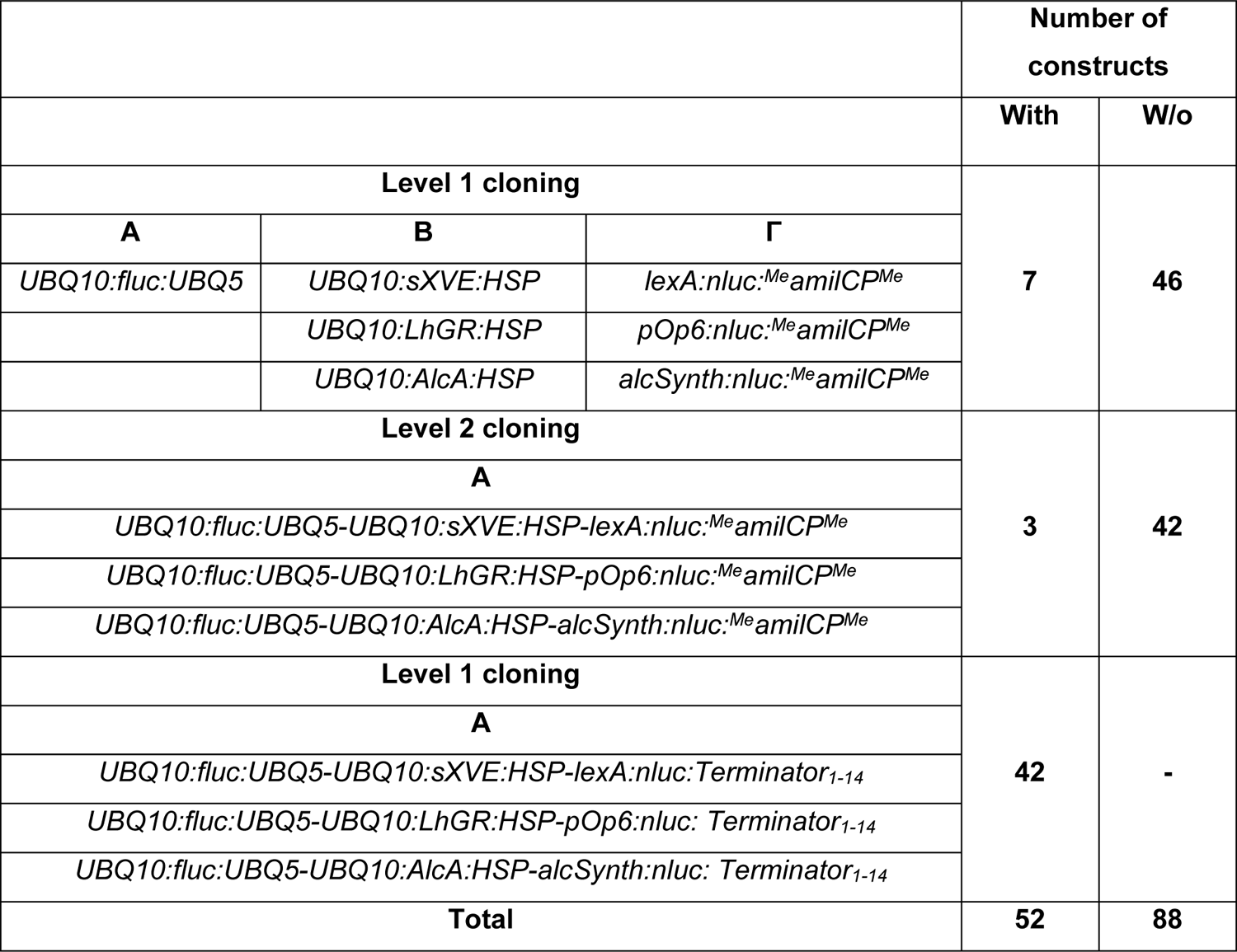
Efficient cloning with MethylAble feature. Application of the MethylAble Feature to generate a library of three inducible systems where the promoters are combined with 14 different terminators. In Level 1 a normaliser unit was built in vector A, the three transactivator units in vector B and three inducible promoters were combined with the MethylAble Feature in vector Γ. Without the MethylAble feature, the inducible promoters should have been combined with all terminators in this step. Then in a Level 2 reaction, the normaliser, the transactivators and the inducible promoters were fused to generate three constructs in total. Lastly, in a level 1 reaction, the inducible constructs were combined with the Level 0 terminators to give the final constructs. Letters represent the Acceptor Vectors.

**Supplementary Table S4.** List of promoter and terminator parts in the standard part library made in this study. 1-10: Genes from which we amplify promoter or/and terminators for likely constitutive plant expression. We selected house-keeping genes, which tend to be ubiquitously expressed, and they are being used as reference genes in qPCR. LEC2 is being expressed in the developing embryo.11-23: Promoters and terminators isolated in other studies and used for characterisation in this work.

| No | TAIR ID | Gene | Protein Function | Pro | Ter | Ref. |
| --- | --- | --- | --- | --- | --- | --- |
| 1 | AT3G18780 | **ACT2**<br>(Actin 2) | Component of the cell cytoskeleton | + | + | This study |
| 2 | AT3G12120 | **FAD2**<br>(Fatty Acid Desaturase 2) | Part of Lipid metabolism | + | + | This study |
| 3 | AT1G27450 | **APT1**<br>(Adenine PhosphoribosylTransferase1) | Contributes to the recycling of adenine | + | + | This study |
| 4 | AT5G18800 | **NDUFA8**<br>(NADH:Ubiquinone oxidoreductase subunit A8) | Mitochondrial electron transport | + | + | This study |
| 5 | AT4G20890 | **TUB9**<br>(Tubulin 9) | A major constituent of microtubules | + | + | This study |
| 6 | AT5G62690 | **TUB2**<br>(Tubulin 2) | A major constituent of microtubules | + | - | This study |
| 7 | AT4G05050 | **UBQ11**<br>(Ubiquitin 11) | Ubiquitin-dependent protein catabolic process | + | - | This study |
| 8 | AT5G20620 | **UBQ4**<br>(Ubiquitin 4) | Ubiquitin-dependent protein catabolic process | + | - | This study |
| 9 | AT1G28300 | **LEC2**<br>(Leafy Cotyledon 1) | Embryo development | + | + | This study |
| 10 | AT5G09810 | **ACT7**<br>(Actin 7) | Component of the cell cytoskeleton | + | - | This study |
| 11 | AT4G05320 | **UBQ10**<br>(Ubiquitin 10) | Ubiquitin-dependent protein catabolic process | + | - | (11) |
| 12 | AT3G16640 | **TCTP**<br>(Translationally Controlled Tumor Protein) | Guanine nucleotide exchange factor in the TOR signalling pathway | + | - | (12) |
| 13 | AT5G38420 | **rbcS-2b**<br>(Rubisco Small Subunit -2b) | Member of the Rubisco small subunit | - | + | (13)<br>Shorter versions in this study |
| 14 | AT5G59720 | **HSP**<br>(Heat shock protein) | Response to heat shock | - | + |  |
| 15 | AT5G59720 | **UBQ5**<br>(Ubiquitin 5) | Ubiquitin-dependent protein catabolic process | - | + |  |
| 16 | Agrobacterium gene | **NOS**<br>(Nopaline synthase) | Nopaline synthesis | + | + | (14) |
| 17 | Agrobacterium gene | **MAS**<br>(Mannopine synthase) | Mannopine Synthesis | + | + |  |
| 18 | Agrobacterium gene | **OCS**<br>(Octopine synthase) | Octopine synthesis | + | - |  |
| 19 | Agrobacterium gene | **G7**<br>(Gene 7) | Unknown function | - | + |  |
| 20 | Cauliflower mosaic virus gene | **35S** | Function in genome replication | + | + | (15) |
| 21 | Synthetic | **G10-90** | 10 G-box sequences fused to CaMV -90/35S promoter | + | - | (16) |
| 22 | Synthetic | **lexA-35S** | lexA operator repeats fused to CaMV 35S core promoter | + | - | (17) |
| 23 | Synthetic | **pOp6-35S** | lacI operator repeats fused to CaMV 35S core promoter | + | - | (18) |
| 24 | Synthetic | **alcSynth-34S** | alcR binding sites fused to FMV 34S core promoter | + | - | (19) |

## REFERENCES

1. Andrianantoandro, E., Basu, S., Karig, D.K. and Weiss, R. (2006) Synthetic biology: new engineering rules for an emerging discipline. Mol. Syst. Biol., 2.

2. Liu, W., Yuan, J.S. and Stewart Jr, C.N. (2013) Advanced genetic tools for plant biotechnology. Nat. Rev. Genet., 14, 781.

3. Naqvi, S., Farré, G., Sanahuja, G., Capell, T., Zhu, C. and Christou, P. (2010) When more is better: multigene engineering in plants. Trends Plant Sci., 15, 48–56.

4. Que, Q., Chilton, M.-D.M., de Fontes, C.M., He, C., Nuccio, M., Zhu, T., Wu, Y., Chen, J.S. and Shi, L. (2010) Trait stacking in transgenic crops: Challenges and opportunities. GM Crops, 1, 220–229.

5. Townson, J. (2017) Recent developments in genome editing for potential use in plants. Biosci. Horizons Int. J. Student Res., 10.

6. Wu, G., Truksa, M., Datla, N., Vrinten, P., Bauer, J., Zank, T., Cirpus, P., Heinz, E. and Qiu, X. (2005) Stepwise engineering to produce high yields of very long-chain polyunsaturated fatty acids in plants. Nat. Biotechnol., 23, 1013–1017.

7. Patron, N.J. (2020) Beyond natural: synthetic expansions of botanical form and function. New Phytol., 227, 295–310.

8. Mortimer, J.C. (2018) Plant synthetic biology could drive a revolution in biofuels and medicine. Exp. Biol. Med., 244, 323–331.

9. Andreou, A.I. and Nakayama, N. (2018) Mobius Assembly: A versatile Golden-Gate framework towards universal DNA assembly. PLoS One, 13, e0189892.

10. Andreou, A.I. and Nakayama, N. (2020) Mobius Assembly BT - DNA Cloning and Assembly: Methods and Protocols. In Chandran, S., George, K.W. (eds). Springer US, New York, NY, pp. 201–218.

11. Weber, E., Engler, C., Gruetzner, R., Werner, S. and Marillonnet, S. (2011) A Modular Cloning System for Standardized Assembly of Multigene Constructs. PLoS One, 6, e16765.

12. Engler, C., Gruetzner, R., Kandzia, R. and Marillonnet, S. (2009) Golden gate shuffling: a one-pot DNA shuffling method based on type IIs restriction enzymes. PLoS One, 4.

13. Ong, J.L., Bilotti, K., Evans Jr, T.C., Potapov, V., Lohman, G.J.S., Langhorst, B.W., Canton, B., Cahoon, D. and Knight, T.F. (2018) A single-molecule sequencing assay for the comprehensive profiling of T4 DNA ligase fidelity and bias during DNA end-joining. Nucleic Acids Res., 46, e79– e79.

14. Potapov, V., Ong, J.L., Kucera, R.B., Langhorst, B.W., Bilotti, K., Pryor, J.M., Cantor, E.J., Canton, B., Knight, T.F., Evans, T.C., et al. (2018) Comprehensive Profiling of Four Base Overhang Ligation Fidelity by T4 DNA Ligase and Application to DNA Assembly. ACS Synth. Biol., 7, 2665–2674.

15. Pryor, J.M., Potapov, V., Kucera, R.B., Bilotti, K., Cantor, E.J. and Lohman, G.J.S. (2020) Enabling one-pot Golden Gate assemblies of unprecedented complexity using data-optimized assembly design. PLoS One, 15, e0238592.

16. Lin, D. and O’Callaghan, C.A. (2018) MetClo: methylase-assisted hierarchical DNA assembly using a single type IIS restriction enzyme. Nucleic Acids Res., 46, e113.

17. Taylor, G.M., Mordaka, P.M. and Heap, J.T. (2018) Start-Stop Assembly: a functionally scarless DNA assembly system optimized for metabolic engineering. Nucleic Acids Res., 47, e17–e17.

18. van Dolleweerd, C.J., Kessans, S.A., Van de Bittner, K.C., Bustamante, L.Y., Bundela, R., Scott, B., Nicholson, M.J. and Parker, E.J. (2018) MIDAS: A Modular DNA Assembly System for Synthetic Biology. ACS Synth. Biol., 7, 1018–1029.

19. Valenzuela-Ortega, M. and French, C. (2019) Joint Universal Modular Plasmids (JUMP): A flexible and comprehensive platform for synthetic biology. bioRxiv, 10.1101/799585.

20. Damalas, S.G., Batianis, C., Martin-Pascual, M., de Lorenzo, V. and Martins dos Santos, V.A.P. (2020) SEVA 3.1: enabling interoperability of DNA assembly among the SEVA, BioBricks and Type IIS restriction enzyme standards. Microb. Biotechnol., **n/a**.

21. Vasudevan, R., Gale, G.A.R., Schiavon, A.A., Puzorjov, A., Malin, J., Gillespie, M.D., Vavitsas, K., Zulkower, V., Wang, B., Howe, C.J., et al. (2019) CyanoGate: A modular cloning suite for engineering cyanobacteria based on the plant MoClo syntax. Plant Physiol., 10.1104/pp.18.01401.

22. Crozet, P., Navarro, F.J., Willmund, F., Mehrshahi, P., Bakowski, K., Lauersen, K.J., Pérez-Pérez, M.-E., Auroy, P., Gorchs Rovira, A., Sauret-Gueto, S., et al. (2018) Birth of a Photosynthetic Chassis: A MoClo Toolkit Enabling Synthetic Biology in the Microalga Chlamydomonas reinhardtii. ACS Synth. Biol., 7, 2074–2086.

23. Garrigues, S., Manzanares, P., Yenush, L., Orzaez, D., Gandía, M., Hernanz-Koers, M. and Marcos, J.F. (2018) FungalBraid: A GoldenBraid-based modular cloning platform for the assembly and exchange of DNA elements tailored to fungal synthetic biology. Fungal Genet. Biol., 116, 51– 61.

24. Marx, H., Egermeier, M. and Sauer, M. (2019) Golden Gate based metabolic engineering strategy for wild-type strains of Yarrowia lipolytica. 10.1093/femsle/fnz022.

25. Pollak, B., Matute, T., Nuñez, I., Cerda, A., Lopez, C., Vargas, V., Kan, A., Bielinski, V., von Dassow, P., Dupont, C.L., et al. (2020) Universal loop assembly: open, efficient and cross-kingdom DNA fabrication. Synth. Biol., 5.

26. Chiasson, D., Giménez-Oya, V., Bircheneder, M., Bachmaier, S., Studtrucker, T., Ryan, J., Sollweck, K., Leonhardt, H., Boshart, M., Dietrich, P., et al. (2019) A unified multi-kingdom Golden Gate cloning platform. Sci. Rep., 9, 10131.

27. Pollak, B., Cerda, A., Delmans, M., Álamos, S., Moyano, T., West, A., Gutiérrez, R.A., Patron, N.J., Federici, F. and Haseloff, J. (2019) Loop assembly: a simple and open system for recursive fabrication of DNA circuits. New Phytol., 222, 628–640.

28. Gantner, J., Ordon, J., Ilse, T., Kretschmer, C., Gruetzner, R., Löfke, C., Dagdas, Y., Bürstenbinder, K., Marillonnet, S. and Stuttmann, J. (2018) Peripheral infrastructure vectors and an extended set of plant parts for the Modular Cloning system. PLoS One, 13, e0197185.

29. Komori, T., Imayama, T., Kato, N., Ishida, Y., Ueki, J. and Komari, T. (2007) Current Status of Binary Vectors and Superbinary Vectors. Plant Physiol., 145, 1155 LP – 1160.

30. Xiang, C., Han, P., Lutziger, I., Wang, K. and Oliver, D.J. (1999) A mini binary vector series for plant transformation. Plant Mol. Biol., 40, 711–717.

31. Coutu, C., Brandle, J., Brown, D., Brown, K., Miki, B., Simmonds, J. and Hegedus, D.D. (2007) pORE: a modular binary vector series suited for both monocot and dicot plant transformation. Transgenic Res., 16, 771–781.

32. Hellens, R.P., Edwards, E.A., Leyland, N.R., Bean, S. and Mullineaux, P.M. (2000) pGreen: a versatile and flexible binary Ti vector for Agrobacterium-mediated plant transformation. Plant Mol. Biol., 42, 819–832.

33. McBride, K.E. and Summerfelt, K.R. (1990) Improved binary vectors for Agrobacterium-mediated plant transformation. Plant Mol. Biol., 14, 269–276.

34. Hamilton, C.M. (1997) A binary-BAC system for plant transformation with high-molecular-weight DNA. Gene, 200, 107–116.

35. Liu, Y.G., Shirano, Y., Fukaki, H., Yanai, Y., Tasaka, M., Tabata, S. and Shibata, D. (1999) Complementation of plant mutants with large genomic DNA fragments by a transformation-competent artificial chromosome vector accelerates positional cloning. Proc. Natl. Acad. Sci. U. S. A., 96, 6535–6540.

36. Pasin, F., Bedoya, L.C., Bernabé-Orts, J.M., Gallo, A., Simón-Mateo, C., Orzaez, D. and García, J.A. (2017) Multiple T-DNA Delivery to Plants Using Novel Mini Binary Vectors with Compatible Replication Origins. ACS Synth. Biol., 6, 1962–1968.

37. Silva, F., Queiroz, J.A. and Domingues, F.C. (2012) Evaluating metabolic stress and plasmid stability in plasmid DNA production by Escherichia coli. Biotechnol. Adv., 30, 691–708.

38. Ertl, P.F. and Thomsen, L.L. (2003) Technical issues in construction of nucleic acid vaccines. Methods, 31, 199–206.

39. Moore, S.J., Lai, H.E., Kelwick, R.J.R., Chee, S.M., Bell, D.J., Polizzi, K.M. and Freemont, P.S. (2016) EcoFlex: A Multifunctional MoClo Kit for E. coli Synthetic Biology. ACS Synth. Biol., 5, 1059–1069.

40. Oliveira, P.H., Prather, K.J., Prazeres, D.M.F. and Monteiro, G.A. (2010) Analysis of DNA repeats in bacterial plasmids reveals the potential for recurrent instability events. Appl. Microbiol. Biotechnol., 87, 2157–2167.

41. Bi, X. and Liu, L.F. (1996) DNA rearrangement mediated by inverted repeats. Proc. Natl. Acad. Sci. U. S. A., 93, 819–823.

42. Lee, S., Su, G., Lasserre, E., Aghazadeh, M.A. and Murai, N. (2012) Small high-yielding binary Ti vectors pLSU with co-directional replicons for Agrobacterium tumefaciens-mediated transformation of higher plants. Plant Sci., 187, 49–58.

43. Jones, H.D., Doherty, A. and Sparks, C.A. (2009) Transient Transformation of Plants BT - Plant Genomics: Methods and Protocols. In Gustafson, J.P., Langridge, P., Somers, D.J. (eds). Humana Press, Totowa, NJ, pp. 131–152.

44. Peremarti, A., Twyman, R.M., Gomez-Galera, S., Naqvi, S., Farre, G., Sabalza, M., Miralpeix, B., Dashevskaya, S., Yuan, D., Ramessar, K., et al. (2010) Promoter diversity in multigene transformation. Plant Mol. Biol., 73, 363–378.

45. Park, S.H., Lee, B.-M., Salas, M.G., Srivatanakul, M. and Smith, R.H. (2000) Shorter T-DNA or additional virulence genes improve Agrobactrium-mediated transformation. Theor. Appl. Genet., 101, 1015–1020.

46. Chung, C.T. and Miller, R.H. (1993) Preparation and storage of competent Escherichia coli cells. Methods Enzymol., 218, 621–627.

47. Huala, E., Dickerman, A.W., Garcia-Hernandez, M., Weems, D., Reiser, L., LaFond, F., Hanley, D., Kiphart, D., Zhuang, M., Huang, W., et al. (2001) The Arabidopsis Information Resource (TAIR): a comprehensive database and web-based information retrieval, analysis, and visualization system for a model plant. Nucleic Acids Res., 29, 102–105.

48. Shahmuradov, I.A., Umarov, R.K. and Solovyev, V. V (2017) TSSPlant: a new tool for prediction of plant Pol II promoters. Nucleic Acids Res., 45, e65–e65.

49. Lescot, M., Dehais, P., Thijs, G., Marchal, K., Moreau, Y., Van de Peer, Y., Rouze, P. and Rombauts, S. (2002) PlantCARE, a database of plant cis-acting regulatory elements and a portal to tools for in silico analysis of promoter sequences. Nucleic Acids Res., 30, 325–327.

50. Higo, K., Ugawa, Y., Iwamoto, M. and Korenaga, T. (1999) Plant cis-acting regulatory DNA elements (PLACE) database: 1999. Nucleic Acids Res., 27, 297–300.

51. Ji, G., Li, L., Li, Q.Q., Wu, X., Fu, J., Chen, G. and Wu, X. (2015) PASPA: a web server for mRNA poly(A) site predictions in plants and algae. Bioinformatics, 31, 1671–1673.

52. Chang, T.-H., Huang, H.-Y., Hsu, J.B.-K., Weng, S.-L., Horng, J.-T. and Huang, H.-D. (2013) An enhanced computational platform for investigating the roles of regulatory RNA and for identifying functional RNA motifs. BMC Bioinformatics, 14, S4.

53. Menges, M. and Murray, J.A.H. (2002) Synchronous Arabidopsis suspension cultures for analysis of cell-cycle gene activity. Plant J., 30, 203–212.

54. Oda, Y. (2017) VND6-induced Xylem Cell Differentiation in Arabidopsis Cell Cultures. Methods Mol. Biol., 1544, 67–73.

55. Chupeau, M.-C., Granier, F., Pichon, O., Renou, J.-P., Gaudin, V. and Chupeau, Y. (2013) Characterization of the early events leading to totipotency in an Arabidopsis protoplast liquid culture by temporal transcript profiling. Plant Cell, 25, 2444–63.

56. Faraco, M., Di Sansebastiano, G. Pietro Spelt, K., Koes, R.E. and Quattrocchio, F.M. (2011) One Protoplast Is Not the Other! Plant Physiol., 156, 474 LP – 478.

57. Mauri, M., Vecchione, S. and Fritz, G. (2019) Deconvolution of Luminescence Cross-Talk in High-Throughput Gene Expression Profiling. ACS Synth. Biol., 8, 1361–1370.

58. Watson, M.R., Lin, Y., Hollwey, E., Dodds, R.E., Meyer, P. and McDowall, K.J. (2016) An Improved Binary Vector and Escherichia coli Strain for Agrobacterium tumefaciens-Mediated Plant Transformation. G3 Genes|Genomes|Genetics, 6, 2195–2201.

59. Bartosik, D., Baj, J., Sochacka, M., Piechucka, E. and Wlodarczyk, M. (2002) Molecular characterization of functional modules of plasmid pWKS1 of Paracoccus pantotrophus DSM 11072. Microbiology, 148, 2847–2856.

60. Challacombe, J.F., Pillai, S. and Kuske, C.R. (2017) Shared features of cryptic plasmids from environmental and pathogenic Francisella species. PLoS One, 12, e0183554.

61. Zaleski, P., Wawrzyniak, P., Sobolewska, A., Łukasiewicz, N., Baran, P., Romańczuk, K., Daniszewska, K., Kierył, P., Płucienniczak, G. and Płucienniczak, A. (2015) pIGWZ12 – A cryptic plasmid with a modular structure. Plasmid, 79, 37–47.

62. Lee, S., Su, G., Lasserre, E., Aghazadeh, M.A. and Murai, N. (2012) Small high-yielding binary Ti vectors pLSU with co-directional replicons for Agrobacterium tumefaciens-mediated transformation of higher plants. Plant Sci., 187, 49–58.

63. Menges, M. and Murray, J.A.H. (2002) Synchronous Arabidopsis suspension cultures for analysis of cell-cycle gene activity. Plant J., 30, 203–212.

64. Naseri, G. and Koffas, M.A.G. (2020) Application of combinatorial optimization strategies in synthetic biology. Nat. Commun., 11, 2446.

65. Guénin, S., Mauriat, M., Pelloux, J., Van Wuytswinkel, O., Bellini, C. and Gutierrez, L. (2009) Normalization of qRT-PCR data: the necessity of adopting a systematic, experimental conditions-specific, validation of references. J. Exp. Bot., 60, 487–493.

66. Harada, J.J. (2001) Role of Arabidopsis LEAFY COTYLEDON genes in seed development. J. Plant Physiol., 158, 405–409.

67. Chupeau, M.-C., Granier, F., Pichon, O., Renou, J.-P., Gaudin, V. and Chupeau, Y. (2013) Characterization of the Early Events Leading to Totipotency in an Arabidopsis Protoplast Liquid Culture by Temporal Transcript Profiling. Plant Cell, 25, 2444–2463.

68. Urquiza-García, U. and Millar, A.J. (2019) Expanding the bioluminescent reporter toolkit for plant science with NanoLUC. Plant Methods, 15, 68.

69. Borghi, L. (2010) Inducible Gene Expression Systems for Plants. Methods Mol. Biol., 655, 65–75.

70. Narsai, R., Howell, K.A., Millar, A.H., O’Toole, N., Small, I. and Whelan, J. (2007) Genome-wide analysis of mRNA decay rates and their determinants in Arabidopsis thaliana. Plant Cell, 19, 3418– 3436.

71. McLauchlan, J., Gaffney, D., Whitton, J.L. and Clements, J.B. (1985) The consensus sequence YGTGTTYY located downstream from the AATAAA signal is required for efficient formation of mRNA 3’ termini. Nucleic Acids Res., 13, 1347–1368.

72. MacNicol, M.C., Cragle, C.E. and MacNicol, A.M. (2011) Context-dependent regulation of Musashi-mediated mRNA translation and cell cycle regulation. Cell Cycle, 10, 39–44.

73. Ghedira, R., De Buck, S., Nolf, J. and Depicker, A. (2013) The efficiency of Arabidopsis thaliana floral dip transformation is determined not only by the Agrobacterium strain used but also by the physiology and the ecotype of the dipped plant. Mol. Plant. Microbe. Interact., 26, 823–832.

74. Wroblewski, T., Tomczak, A. and Michelmore, R. (2005) Optimization of Agrobacterium-mediated transient assays of gene expression in lettuce, tomato and Arabidopsis. Plant Biotechnol. J., 3, 259–273.

75. Wu, H.-Y., Liu, K.-H., Wang, Y.-C., Wu, J.-F., Chiu, W.-L., Chen, C.-Y., Wu, S.-H., Sheen, J. and Lai, E.-M. (2014) AGROBEST: an efficient Agrobacterium-mediated transient expression method for versatile gene function analyses in Arabidopsis seedlings. Plant Methods, 10, 19.

76. Li, X.-Q. and Du, D. (2014) Motif types, motif locations and base composition patterns around the RNA polyadenylation site in microorganisms, plants and animals. BMC Evol. Biol., 14, 162.

77. Han, Y.-J., Kim, Y.-M., Hwang, O.-J. and Kim, J.-I. (2015) Characterization of a small constitutive promoter from Arabidopsis translationally controlled tumor protein (AtTCTP) gene for plant transformation. Plant Cell Rep., 34, 265–275.

78. Jiang, P., Zhang, K., Ding, Z., He, Q., Li, W., Zhu, S., Cheng, W., Zhang, K. and Li, K. (2018) Characterization of a strong and constitutive promoter from the Arabidopsis serine carboxypeptidase-like gene AtSCPL30 as a potential tool for crop transgenic breeding. BMC Biotechnol., 18, 59.

79. Cai, Y.-M., Kallam, K., Tidd, H., Gendarini, G., Salzman, A. and Patron, N.J. (2020) Rational design of minimal synthetic promoters for plants. Nucleic Acids Res., 10.1093/nar/gkaa682.

80. Engler, C., Youles, M., Gruetzner, R., Ehnert, T.M., Werner, S., Jones, J.D.G., Patron, N.J. and Marillonnet, S. (2014) A Golden Gate modular cloning toolbox for plants. ACS Synth. Biol., 3, 839– 843.

81. Sarrion-Perdigones, A., Vazquez-Vilar, M., Palaci, J., Castelijns, B., Forment, J., Ziarsolo, P., Blanca, J., Granell, A. and Orzaez, D. (2013) GoldenBraid 2.0: A Comprehensive DNA Assembly Framework for Plant Synthetic Biology. Plant Physiol., 162, 1618–1631.

82. Yanagisawa, S. (2000) Dof1 and Dof2 transcription factors are associated with expression of multiple genes involved in carbon metabolism in maize. Plant J., 21, 281–288.

83. Biłas, R., Szafran, K., Hnatuszko-Konka, K. and Kononowicz, A.K. (2016) Cis-regulatory elements used to control gene expression in plants. Plant Cell. Tissue Organ Cult., 127, 269–287.

84. Schlücking, K., Edel, K.H., Köster, P., Drerup, M.M., Eckert, C., Steinhorst, L., Waadt, R., Batistič, O. and Kudla, J. (2013) A New β-Estradiol-Inducible Vector Set that Facilitates Easy Construction and Efficient Expression of Transgenes Reveals CBL3-Dependent Cytoplasm to Tonoplast Translocation of CIPK5. Mol. Plant, 6, 1814–1829.

85. Craft, J., Samalova, M., Baroux, C., Townley, H., Martinez, A., Jepson, I., Tsiantis, M. and Moore, I. (2005) New pOp/LhG4 vectors for stringent glucocorticoid-dependent transgene expression in Arabidopsis. Plant J., 41, 899–918.

86. Ansari, A. and Hampsey, M. (2005) A role for the CPF 3′ -end processing machinery in RNAP II-dependent gene looping. Genes Dev., 19, 2969–2978.

87. Calvo, O. and Manley, J.L. (2003) Strange bedfellows: Polyadenylation factors at the promoter. Genes Dev., 17, 1321–1327.

88. Hampsey, M., Singh, B.N., Ansari, A., Lainé, J.P. and Krishnamurthy, S. (2011) Control of eukaryotic gene expression: Gene loops and transcriptional memory. Adv. Enzyme Regul., 10.1016/j.advenzreg.2010.10.001.

89. Moabbi, A.M., Agarwal, N., El Kaderi, B. and Ansari, A. (2012) Role for gene looping in intron-mediated enhancement of transcription. Proc. Natl. Acad. Sci., 109, 8505 LP – 8510.

90. Tan-Wong, S.M., Zaugg, J.B., Camblong, J., Xu, Z., Zhang, D.W., Mischo, H.E., Ansari, A.Z., Luscombe, N.M., Steinmetz, L.M. and Proudfoot, N.J. (2012) Gene loops enhance transcriptional directionality. Science, 338, 671–675.

91. Al Husini, N., Kudla, P. and Ansari, A. (2013) A Role for CF1A 3′ End Processing Complex in Promoter-Associated Transcription. PLOS Genet., 9, e1003722.

92. Mukundan, B. and Ansari, A. (2013) Srb5/Med18-mediated termination of transcription is dependent on gene looping. J. Biol. Chem., 288, 11384–11394.

93. Medler, S. and Ansari, A. (2015) Gene looping facilitates TFIIH kinase-mediated termination of transcription. Sci. Rep., 5, 12586.

94. Pérez-González, A. and Caro, E. (2018) Effect of transcription terminator usage on the establishment of transgene transcriptional gene silencing. BMC Res. Notes, 11, 511.

95. Beyene, G., Buenrostro-Nava, M.T., Damaj, M.B., Gao, S.J., Molina, J. and Mirkov, E.E. (2011) Unprecedented enhancement of transient gene expression from minimal cassettes using a double terminator. Plant Cell Rep., 30, 13–25.

96. Yamamoto, T., Hoshikawa, K., Ezura, K., Okazawa, R., Fujita, S., Takaoka, M., Mason, H.S., Ezura, H. and Miura, K. (2018) Improvement of the transient expression system for production of recombinant proteins in plants. Sci. Rep., 8, 4755.

97. Diamos, A.G. and Mason, H.S. (2018) Chimeric 3’ flanking regions strongly enhance gene expression in plants. Plant Biotechnol. J., 16, 1971–1982.

## Supplementary References

1. Moore, S.J., Lai, H.E., Kelwick, R.J.R., Chee, S.M., Bell, D. J., Polizzi, K.M. and Freemont, P.S. (2016) EcoFlex: A Multifunctional MoClo Kit for E. coli Synthetic Biology. ACS Synth. Biol., 5, 1059–1069.

2. Lazo, G.R., Stein, P.A. and Ludwig, R.A. (1991) A DNA Transformation–Competent Arabidopsis Genomic Library in Agrobacterium. Bio/Technology, 9, 963–967.

3. Tao, L., Jackson, R.E. and Cheng, Q. (2005) Directed evolution of copy number of a broad host range plasmid for metabolic engineering. Metab. Eng., 7, 10–17.

4. Wroblewski, T., Tomczak, A. and Michelmore, R. (2005) Optimization of Agrobacterium-mediated transient assays of gene expression in lettuce, tomato and Arabidopsis. Plant Biotechnol. J., 3, 259–273.

5. Slater, S.C., Goodner, B.W., Setubal, J.C., Goldman, B.S., Wood, D.W. and Nester, E.W. (2008) The Agrobacterium Tumefaciens C58 Genome BT - Agrobacterium: From Biology to Biotechnology. In Tzfira, T., Citovsky, V. (eds). Springer New York, New York, NY, pp. 149–181.

6. Van Larebeke, N., Engler, G., Holsters, M., Van Den Elsacker, S., Zaenen, I., Schilperoort, R.A. and Schell, J. (1974) Large plasmid in agrobacterium tumefaciens essential for crown gall-inducing ability. Nature, 252, 169–170.

7. Koncz, C. and Schell, J. (1986) The promoter of TL-DNA gene 5 controls the tissue-specific expression of chimaeric genes carried by a novel type of Agrobacterium binary vector. Mol. Gen. Genet. MGG, 204, 383–396.

8. Van Larebeke, N., Engler, G., Holsters, M., Van Den Elsacker, S., Zaenen, I., Schilperoort, R.A. and Schell, J. (1974) Large plasmid in agrobacterium tumefaciens essential for crown gall-inducing ability. Nature, 252, 169–170.

9. Hamilton, C.M. (1997) A binary-BAC system for plant transformation with high-molecular-weight DNA. Gene, 200, 107–116.

10. Hoekema, A., Hirsch, P.R., Hooykaas, P.J.J. and Schilperoort, R.A. (1983) A binary plant vector strategy based on separation of vir- and T-region of the Agrobacterium tumefaciens Ti-plasmid. Nature, 303, 179–180.

11. Grefen, C., Donald, N., Hashimoto, K., Kudla, J., Schumacher, K. and Blatt, M.R. (2010) A ubiquitin-10 promoter-based vector set for fluorescent protein tagging facilitates temporal stability and native protein distribution in transient and stable expression studies. Plant J., 64, 355–365.

12. Han, Y.-J., Kim, Y.-M., Hwang, O.-J. and Kim, J.-I. (2015) Characterization of a small constitutive promoter from Arabidopsis translationally controlled tumor protein (AtTCTP) gene for plant transformation. Plant Cell Rep., 34, 265–275.

13. Nagaya, S., Kawamura, K., Shinmyo, A. and Kato, K. (2010) The HSP terminator of arabidopsis thaliana increases gene expression in plant cells. Plant Cell Physiol., 51, 328–332.

14. Engler, C., Youles, M., Gruetzner, R., Ehnert, T.M., Werner, S., Jones, J.D.G., Patron, N.J. and Marillonnet, S. (2014) A Golden Gate modular cloning toolbox for plants. ACS Synth. Biol., 3, 839– 843.

15. Benfey, P.N. and Chua, N.H. (1990) The Cauliflower Mosaic Virus 35S Promoter: Combinatorial Regulation of Transcription in Plants. Science, 250, 959–966.

16. Ishige, F., Takaichi, M., Foster, R., Chua, N.-H. and Oeda, K. (1999) A G-box motif (GCCACGTGCC) tetramer confers high-level constitutive expression in dicot and monocot plants. Plant J., 18, 443– 448.

17. Schlücking, K., Edel, K.H., Köster, P., Drerup, M.M., Eckert, C., Steinhorst, L., Waadt, R., Batistič, O. and Kudla, J. (2013) A New β-Estradiol-Inducible Vector Set that Facilitates Easy Construction and Efficient Expression of Transgenes Reveals CBL3-Dependent Cytoplasm to Tonoplast Translocation of CIPK5. Mol. Plant, 6, 1814–1829.

18. Craft, J., Samalova, M., Baroux, C., Townley, H., Martinez, A., Jepson, I., Tsiantis, M. and Moore, I. (2005) New pOp/LhG4 vectors for stringent glucocorticoid-dependent transgene expression in Arabidopsis. Plant J., 41, 899–918.

19. Pasin, F., Bedoya, L.C., Bernabé-Orts, J.M., Gallo, A., Simón-Mateo, C., Orzaez, D. and García, J.A. (2017) Multiple T-DNA Delivery to Plants Using Novel Mini Binary Vectors with Compatible Replication Origins. ACS Synth. Biol., 6, 1962–1968.

